# Multi-omic profiling identifies malignant subpopulations and a hypoxia-driven angiogenic axis as therapeutic vulnerabilities in sinonasal squamous cell carcinoma

**DOI:** 10.1101/2024.12.01.626270

**Authors:** Chaelin You, Jaewoo Park, Jung Yeon Jang, Myeong Sang Yu, Yoo-Sam Chung, Keunsoo Kang, Jihwan Park, Ji Heui Kim, Kyuho Kang

**Author notes:** Correspondence: Kyuho Kang,; Ji Heui Kim.

## Abstract

Sinonasal squamous cell carcinoma (SNSCC) is a rare and aggressive malignancy with limited treatment options, necessitating comprehensive molecular characterization to uncover actionable therapeutic targets. Here, we integrate bulk and single-cell transcriptomics, epigenomics, and DNA methylation profiling to construct a detailed molecular and cellular landscape of SNSCC. Our analysis identifies five transcriptionally and epigenetically distinct malignant subpopulations, including hypoxic (TC1) and proliferative (TC2) clusters, with TC1 showing significant association with poor clinical outcomes. We uncover a hypoxia-driven angiogenic axis, wherein TC1 cells secrete adrenomedullin (ADM), MIF, and VEGFA to activate endothelial tip cells, fostering tumor angiogenesis. Epigenetic reprogramming, characterized by alterations in DNA methylation and chromatin accessibility, underpins these transcriptional programs, with AP-1 transcription factors emerging as key regulators. Notably, ADM expression is epigenetically controlled, correlates with advanced clinical stage, and predicts reduced survival in head and neck cancer patients. This multi-omic study highlights the tumor heterogeneity and microenvironmental interactions driving SNSCC progression, revealing epigenetically regulated pathways as promising therapeutic targets for future intervention.

## Introduction

Sinonasal cancer is a rare subset of head and neck cancers, comprising less than 1% of all malignant neoplasms and 5% of head and neck malignancies^1, 2, 3^, and is characterized by highly heterogeneous and distinct clinicopathological features that distinguish it from other head and neck malignancies^4, 5^. Sinonasal squamous cell carcinoma (SNSCC), the predominant histological subtype^5, 6^, primarily affects the maxillary sinuses (60%) and nasal cavity (25%)^1, 7^. Late-stage diagnosis is common, often involving surrounding bone invasion and resulting in poor prognosis^8, 9, 10^. Despite advances in cancer therapeutics, the 5-year survival rate for SNSCC patients remains low, ranging from 30-50%^9, 11^, with little improvement observed in recent decades^1, 8^. The rarity of SNSCC poses significant challenges, including limited sample availability for research, diagnostic complexities, and restricted treatment options^12, 13^. Current therapeutic approaches are largely adapted from protocols designed for more common malignancies^4, 5^, highlighting an urgent need for targeted research to understand disease mechanisms and identify novel therapeutic approaches.

Recent technological advancements in transcriptomics and epigenomics have significantly enhanced our understanding of cancer biology, including rare malignancies^14^. Bulk RNA sequencing (RNA-seq) has elucidated gene expression profiles in complex tumor tissues across various cancers^15^, while comprehensive epigenetic profiling has revealed regulatory mechanisms related to oncogenic gene transcription in pan-cancer studies^16, 17^. In SNSCC, transcriptome analyses have identified potential biomarkers and delineated comparative features of subtypes^18, 19, 20, 21^. Additionally, large-scale DNA methylation profiling has uncovered distinct subtypes of sinonasal cancer, particularly in undifferentiated carcinomas, improving diagnostic efficacy and highlighting the importance of epigenetic regulation^22^.

The tumor microenvironment (TME) plays a crucial role in cancer progression and treatment response^23, 24^. Single-cell technologies, including single-cell RNA sequencing (scRNA-seq) and epigenomic profiling, have provided unprecedented insights into TME components and characteristics by revealing cellular heterogeneity and diverse phenotypes within tumors^25, 26^. Multi-omic approaches, combining gene expression and chromatin accessibility analyses at the single-cell level, have offered more concrete evidence of cell identity and properties in various cancers^27, 28, 29^. A recent investigation into a rare cancer, desmoplastic small round cell tumor (DSRCT), has integrated single-cell multi-omics approaches to uncover heterogeneous features of intra- and inter-tumoral landscapes, as well as gene signatures associated with clinical outcomes^30^. While these single-cell approaches have yielded valuable insights into other types of head and neck cancers^31, 32, 33^, they have not yet been applied to sinonasal cancers. The intricate cellular and molecular landscapes of the TME in SNSCC, including its cellular composition and intercellular communication networks, remain poorly defined, necessitating comprehensive profiling at the single-cell level.

Hypoxia represents a particularly critical aspect of the TME in SNSCC that warrants investigation. This common feature of solid tumors drives aggressive phenotypes and treatment resistance^34, 35^. While hypoxic signatures have been reported in head and neck cancers^36^, their specific relevance to SNSCC pathogenesis remains poorly understood. Previous studies have demonstrated oxygen-deprived features in SNSCC. Hypoxia-associated proteins, including CA-IX (positive in 85% of cases), VEGF (85% of cases), VEGF-R1 (75% of cases), and GLUT-1 (62.5% of cases), were overexpressed in SNSCC patients^37^, underscoring the significance of hypoxia in this malignancy. Furthermore, larger tumors displayed more heterogeneous characteristics with regions of necrosis^38^, which develops in response to chronic hypoxic conditions^39^. Further investigation of hypoxia’s impact on transcriptional regulation and cell-cell interactions in SNSCC could reveal novel therapeutic targets.

In this study, we present an integrated multi-omic profiling analysis to characterize the molecular features and TME of SNSCC. Our dataset encompasses bulk RNA sequencing, DNA methylation arrays, and matched single-nucleus RNA and ATAC sequencing (snRNA/ATAC-seq) from SNSCC tumors and normal nasal mucosa. We identify core oncogenic programs, epigenetic regulatory mechanisms, and heterogeneous cellular populations driving SNSCC pathogenesis. Our analyses reveal distinct malignant cell subsets with prognostic relevance and uncover a hypoxia-angiogenesis axis mediated by specific tumor-endothelial cell interactions. These findings provide a high-resolution map of the SNSCC cellular ecosystem and highlight potential therapeutic vulnerabilities in this challenging malignancy.

## Results

### Transcriptome analysis identifies distinct gene signatures and characteristics of SNSCC

To comprehensively characterize SNSCC, we investigated transcriptomic and epigenomic alterations in tumor and normal tissues obtained from patients, along with nasal mucosa tissues from healthy donors, at both bulk and single-nucleus levels (Fig. 1A and Table 1). We performed bulk RNA sequencing (RNA-seq) on surgically resected specimens from eight patients with SNSCC including seven tumor and seven normal tissues. All tumor samples were diagnosed as stage IV (T4a or T4b) HPV-negative sinonasal carcinoma, with paired normal tissue collected from all patients except one. Detailed clinical information of patients is shown in Table 2. We clearly segregated tumor and normal tissues based on principal component analysis (PCA) and identified 806 differentially expressed genes (DEGs) including 442 upregulated and 364 downregulated genes in tumors relative to normal tissues (adjusted *p*-value < 0.05, > 2-fold differences in expression) (Fig. 1B and Supplementary Fig. 1A).

**Fig. 1.**
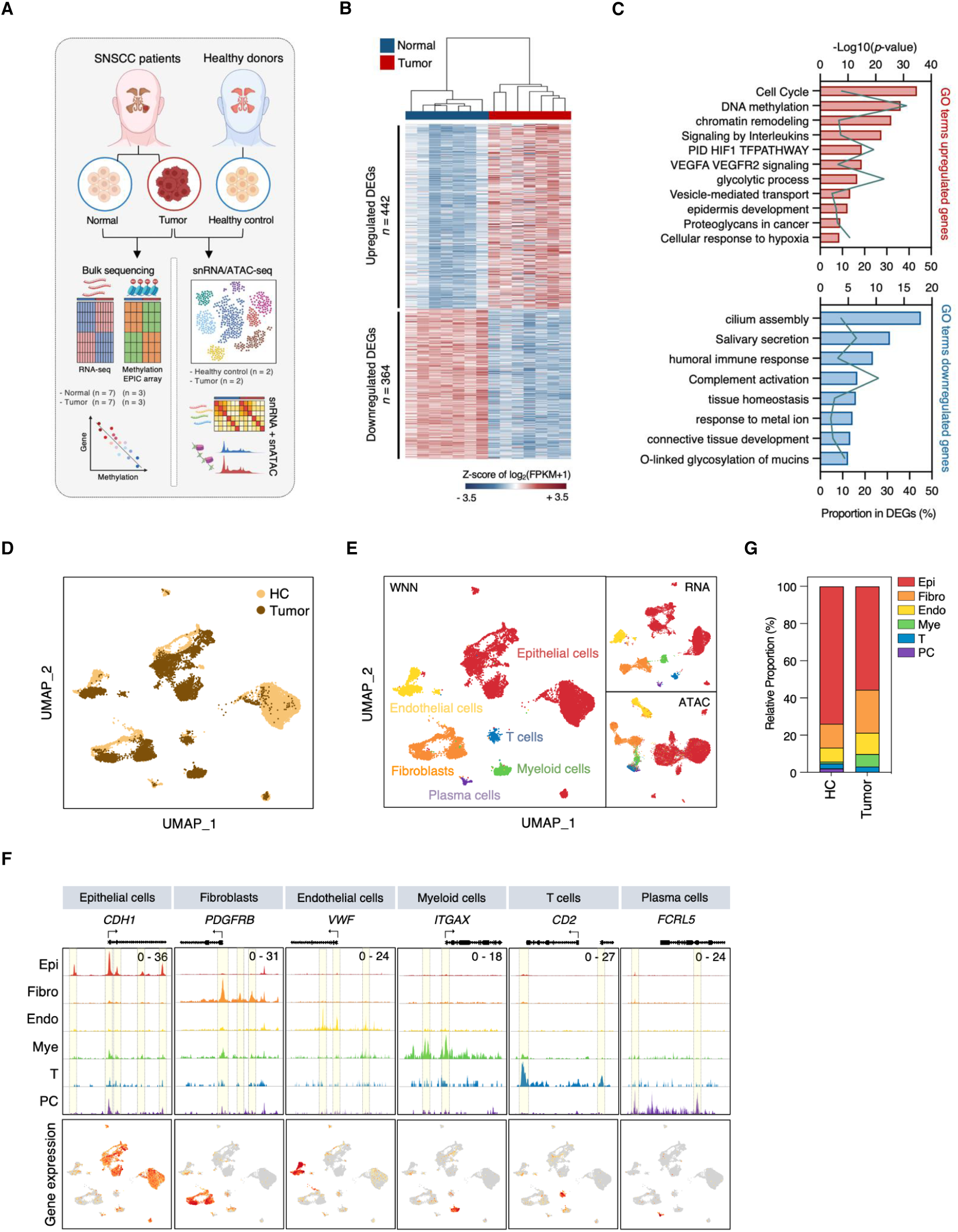
Global transcriptomic characteristics and cellular diversity in SNSCC. **A** Overview of sinonasal squamous cell carcinoma (SNSCC) bulk and single-cell profiling study. Created with Biorender.com. **B** K-means (k = 2) clustering of all differentially expressed genes (DEGs) between tumor and normal tissues from SNSCC patients. A gene is considered significant ifs fold change is > 2, adjusted *p*-value < 0.05, and fragments per kilobase of transcript per million mapped reads (FPKM) > 2. Expression values (log_2_(FPKM+1)) are Z score transformed. **C** Gene ontology (GO) enrichment analysis of DEGs between tumor and normal tissues from SNSCC patients using the METASCAPE online tool. **D** Uniform manifold approximation and projection (UMAP) visualization of 17,648 nuclei from 2 SNSCC and 2 healthy control (HC) tissues colored by tissue type. **E** UMAP visualization of the weighted nearest neighbor (WNN) clustering by WNN, RNA, or ATAC profiles colored by cell type. **F** Proportion of major cell types in SNSCC versus HC tissues. **G** Chromatin accessibility (upper) and gene expression (lower) of cell type marker genes across major cell types. Peaks showing differences between cell types are highlighted.

**Table 1.**
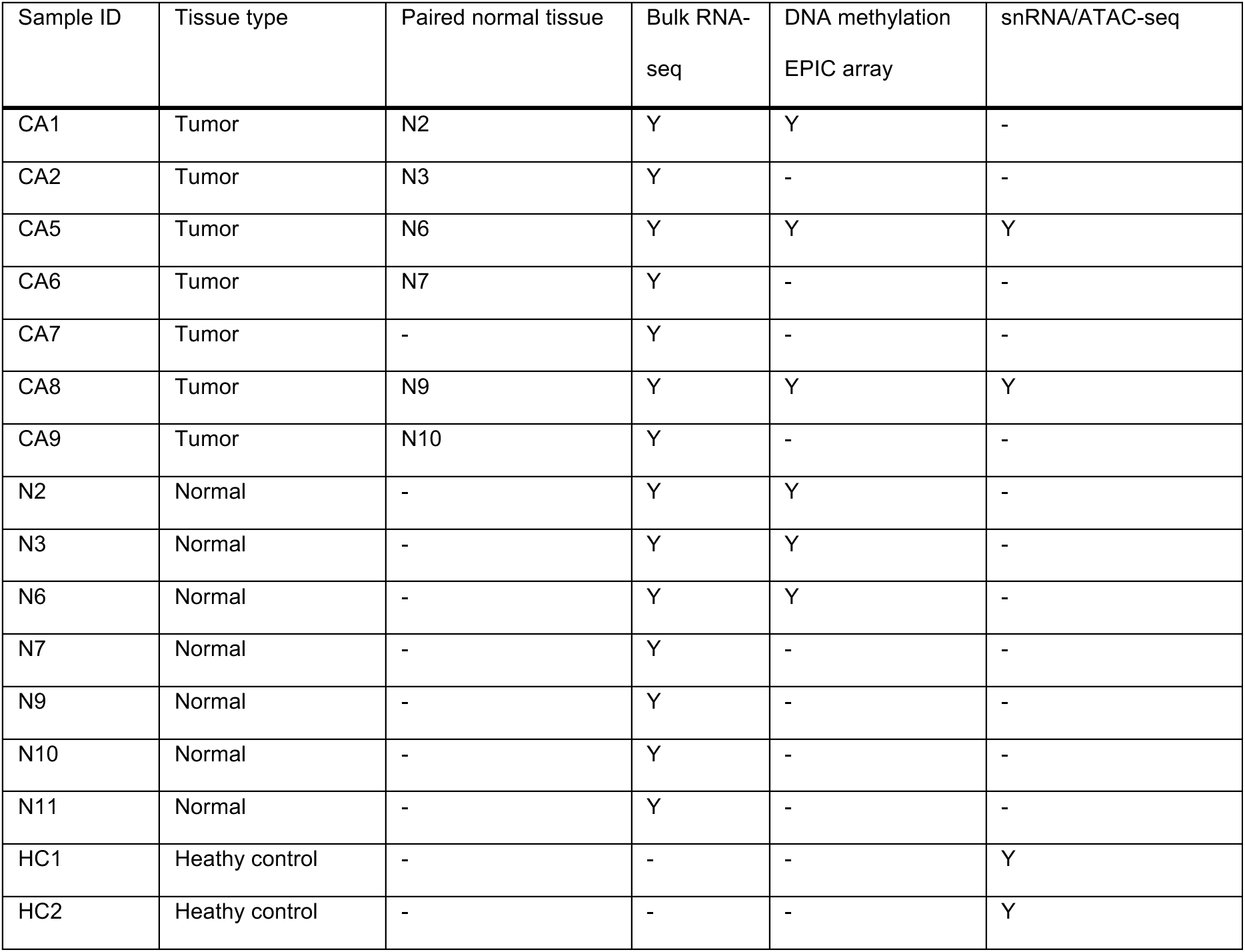
Summary of sequencing information

**Table 2.** Clinical data for SNSCC patients

| Patient ID | Sex | Age | Subsite | Primary | T | Histology | IP<sup>d</sup>_Inverted | Tumor grade | Lymphovascular invasion | Perineural invasion |
| --- | --- | --- | --- | --- | --- | --- | --- | --- | --- | --- |
| CA1 | M | 71 | FS<sup>a</sup> | Primary | 4b | SCC | IP background | 3 | Neg | Pos |
| CA2 | M | 48 | MS<sup>b</sup> | Primary | 4a | SCC | - | 2 | N/A | Pos |
| CA5 | M | 75 | NC<sup>c</sup> | Primary | 4a | SCC | - | 2 | Pos | Neg |
| CA6 | M | 73 | FS | Rec | 4b | SCC | - | 2 | Neg | Neg |
| CA7 | F | 70 | FS | Rec | 4b | SCC c IP | IP background | 1 | N/A | N/A |
| CA8 | M | 61 | NC | Primary | 4a | SCC | IP background | 2 | Pos | Neg |
| CA9 | M | 78 | MS | Primary | 4a | SCC | - | 2 | Neg | Neg |

<sup>a</sup>Frontal sinus; <sup>b</sup>Maxillary sinus; <sup>c</sup>Nasal cavity; <sup>d</sup>Inverted papilloma

Gene ontology (GO) analysis revealed distinct molecular functions in SNSCC. Upregulated genes were notably enriched in cell cycle regulation (*MKI67* and *TOP2A*) (Fig. 1C). GO terms associated with epigenetic regulation, including DNA methylation and chromatin remodeling, were overrepresented, with critical regulators of chromatin structure such as *DNMT1* (DNA methyltransferase) and *KDM3A* (histone demethylase) showing increased expression in SNSCC. In relation to metabolic processes, the HIF1 TF pathway, glycolytic process, and cellular response to hypoxia were enriched, with key mediators such as *HIF1A* and *SLC2A1* (also known as *GLUT1*) considerably elevated in SNSCC (Fig. 1C and Supplementary Fig. 1B). By contrast, downregulated genes were related to cilium assembly, salivary secretion, and O-linked glycosylation of mucins, reflecting typical characteristics of epithelial layers within mucosal tissues. For instance, genes involved in mucin secretion (*MUC5AC* and *MUC5B*), which are essential for airway epithelium defense mechanisms^40^, were significantly reduced in tumors (Fig. 1C and Supplementary Fig. 1B). Gene set enrichment analysis (GSEA) using the hallmark gene sets confirmed these findings, revealing E2F targets, G2M checkpoint, hypoxia, and glycolysis as remarkably enriched cellular processes in overexpressed genes (Supplementary Fig. 1C).

The transcriptome analysis also revealed distinct immune-related signatures. Signaling by interleukins was enriched among upregulated genes, whereas humoral immune response and complement activation were enriched among downregulated genes. Consistently, immunoregulatory genes such as *CCL3* and *SPP1*, which regulate immune cell infiltration, were considerably enhanced, while genes associated with humoral responses (*DMBT1* and *XBP1*) and complement components (*C3* and *C4A*) were significantly repressed (Fig. 1C and Supplementary Fig. 1B). These results demonstrate that SNSCC exhibits a distinct molecular profile characterized by dysregulated cell cycle control, epigenetic modifications, metabolic reprogramming, and immune responses that fundamentally distinguish tumor tissue from its normal counterpart.

### Single-cell profiling unveils cellular composition and diversity in SNSCC

To gain a deeper understanding of the TME in SNSCC, we conducted combined snRNA/ATAC-seq analysis of tumor tissues from patients with SNSCC (*n* = 2) and healthy nasal mucosa tissues (*n* = 2; hereafter ’healthy control (HC)’) using the 10x Genomics Multiome platform, which enables mapping of gene expression and chromatin accessibility in individual nuclei (Fig. 1A). After quality control and Harmony batch correction, we obtained a total of 18,319 cells. Using weighted nearest neighbor (WNN) analysis to combine RNA and ATAC profiles, we identified 28 clusters comprising six major cell types (Fig. 1D, E and Supplementary Fig. 1D). Undefined clusters labeled as ’Other’, which were either positive for multiple marker genes or consisted of low-quality cells, were excluded from further analysis (Supplementary Fig. 1E, F). Each cell compartment was supported by both gene expression and open chromatin states of representative known cell type markers (Fig. 1F and Supplementary Fig. 1E), including epithelial cells (*CDH1*, *EPCAM*, and *KRT8*), fibroblasts (*DCN*, *PDGFRB*, and *THY1*), endothelial cells (*CD34*, *CDH5*, and *VWF*), myeloid cells (*ITGAX*, *MS4A6A*, and *SPI1*), T cells (*CD2*, *CD3D*, and *CD3G*), and plasma cells (*FCRL5*, *JCHAIN*, and *MZB1*).

The relative proportions of the six major cell types in tumor and HC tissues are shown in Fig. 1G. Epithelial cell populations were predominant in both tumor and HC tissues. Compared to HCs, tumors showed an increased proportion of myeloid cells and a decreased proportion of plasma cells (Fig. 1G and Supplementary Fig. 1G). To further evaluate immune cell infiltration between normal and tumor tissues from SNSCC patients, we estimated the fractions of 22 immune cell types in bulk RNA-seq data using CIBERSORTx. Consistent with our snRNA/ATAC-seq findings, macrophages were significantly increased, whereas plasma cells were decreased in SNSCC (Supplementary Fig. 1H, I). Additionally, cell differentiation markers including *CD68* (a pan-macrophage marker) and *JCHAIN* (a plasma cell marker) were detected among the upregulated and downregulated genes, respectively (Supplementary Fig. 1J).

### Cell type-specific transcriptome analysis highlight oncogenic functions in SNSCC

To better characterize cell type-specific gene expression profiles between tumor and HC tissues, we identified DEGs across various cell types that revealed distinct transcriptional programs between tumor and normal tissue environments (adjusted *p*-value < 0.05, > 1 log_2_-fold change) (Fig. 2A). Due to the low abundance of plasma cells (< 20 cells) in tumor tissues, only a small number of DEGs were detected in this population. Analysis of distinct cellular functions in tumor-specific cell types uncovered key pathways associated with SNSCC (Fig. 2B, C and Supplementary Fig. 2A). Epithelial cells in SNSCC were characterized by upregulation of hypoxia- and glycolysis-related genes (*ADM*, *ENO1*, *LDHA*, *NDRG1*, *PKM*, and *SLC2A1*) and epithelial development-related genes (*CDH3*, *DSC3*, *DSG3*, *KRT6A*, *KRT17*, and *SFN*) (Fig. 2B), corroborating our bulk RNA-seq findings. Tumor fibroblasts prominently expressed known cancer-associated fibroblast (CAF) markers such as *FAP* and *POSTN*, along with genes associated with collagen formation (*COL1A1*, *COL1A2*, and *COL6A1*) and TGFβ signaling (*TGFB2*, *TGFBR1*, *ITGAV*, and *ITGB1*) (Fig. 2B). Tumor endothelial cells showed enrichment in angiogenesis-related GO terms, including VEGFA-VEGFR2 signaling, sprouting angiogenesis, and regulation of vascular permeability (Supplementary Fig. 2A). Notably, genes associated with angiogenic sprouts (*ACVRL1*, *ANGPT2*, *FLT4*, and *KDR*) and NOTCH receptors (*NOTCH1* and *NOTCH4*), crucial for angiogenic growth^41^, were upregulated in tumor endothelial cells, indicating active angiogenesis in SNSCC patients (Fig. 2B).

**Fig. 2.**
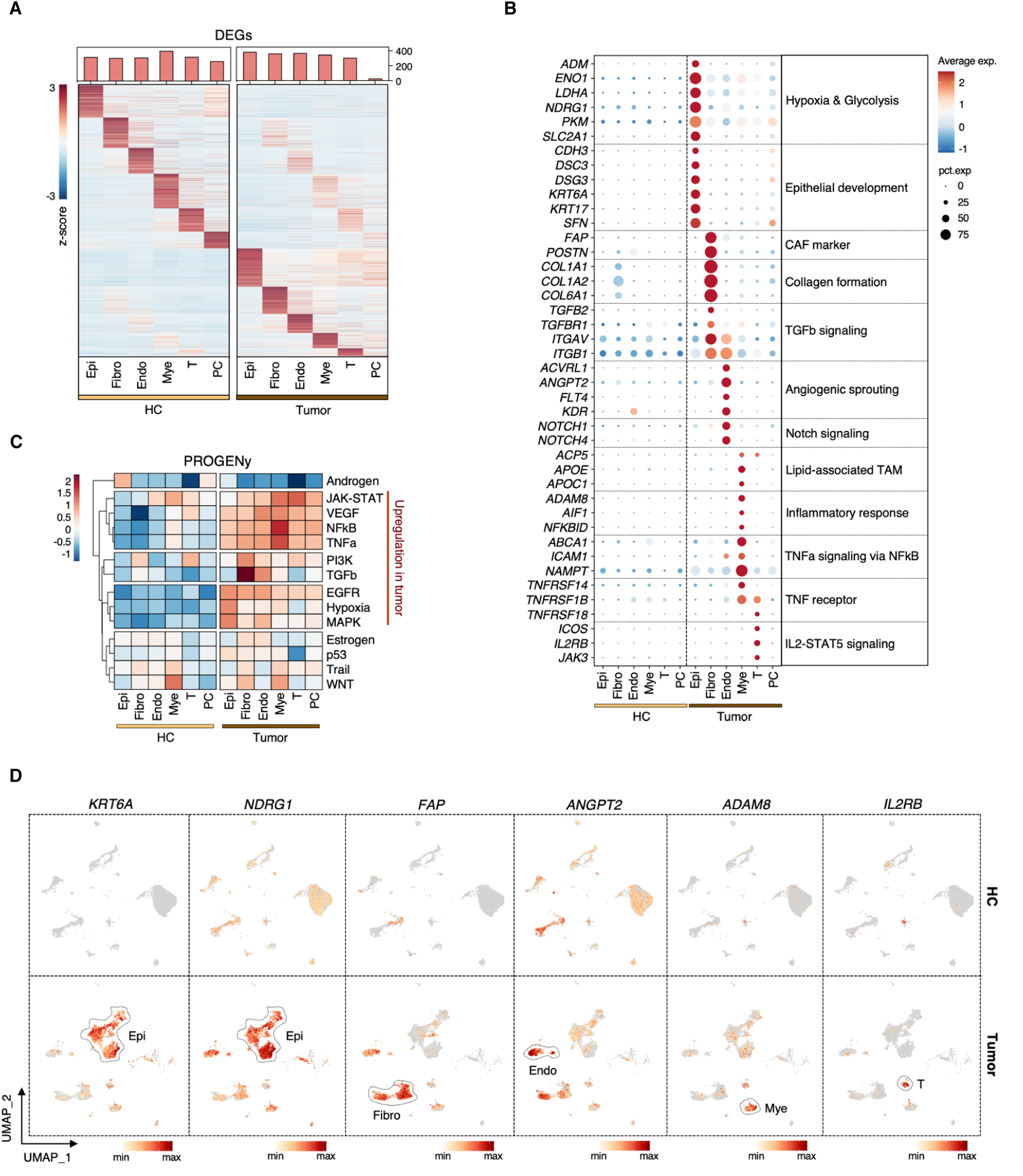
Cell type-specific single-cell transcriptome landscapes and functional analysis in SNSCC. **A** Heatmap of DEGs in each cell type within SNSCC and HC tissues. **B** Dot plot illustrating the functional characteristics and associated genes for each cell type in SNSCC. **C** Heatmap showing the activity of 14 signaling pathways determined using PROGENy, comparing cell types in SNSCC and HC tissues. **D** Feature plots displaying the expression of *KRT6A*, *NDRG1*, *FAP*, *ANGPT2*, *ADAM8*, and *IL2RB* identified in **B** across all cell types in SNSCC tumor and HC tissues.

Chronic inflammation induced by mucosal irritants such as organic dust can contribute to sinonasal cancer development^5^. For instance, organic dust exposure can stimulate macrophages to produce inflammatory cytokines and chemokines, including TNF, which may facilitate tumorigenesis^42, 43^. We found that myeloid cells, the predominant immune compartment in SNSCC (Fig. 1G), exhibited complex biological features, including lipid-associated tumor-associated macrophage (TAM) signatures^44^ (*ACP5*, *APOE*, and *APOC1*) and inflammatory responses (*ADAM8*, *AIF1* and *NFKBID*). Additionally, genes involved in TNF signaling *via* NFκB (*ABCA1*, *ICAM1*, and *NAMPT*) were significantly upregulated in tumor myeloid cells (Fig. 2B). Tumor-infiltrating T cells showed enrichment for primary immunodeficiency and JAK-STAT signaling pathway ontologies, with enhanced expression of TNF receptors (*TNFRSF1B* and *TNFRSF18*) and IL2-STAT5 signaling-associated genes (*ICOS*, *IL2RB*, and *JAK3*) (Fig. 2B and Supplementary Fig. 2A), indicating that immune responses in SNSCC are primarily mediated by myeloid and T cells.

GO pathways in healthy nasal mucosa reflected cell type-specific characteristics (Supplementary Fig. 2B). Endothelial cells showed enriched vascular networks, myeloid cells exhibited phagocytosis, and T cells displayed T cell receptor signaling activation, paralleling observations in SNSCC. HC epithelial cells were enriched for salivary secretion-related genes (*CCL28*, *DMBT1*, *EGF*, and *STATH*), consistent with our bulk RNA-seq findings (Supplementary Fig. 2B, C) and reflecting the airway’s ciliated and secretory cell composition^40^. Normal fibroblasts showed higher expression of muscle structure-related genes (*ACTA2*, *FHL1*, *MYH11*, and *MYOCD*) compared to CAFs (Supplementary Fig. 2B, C).

PROGENy pathway analysis revealed elevated pathway activity scores for JAK-STAT, VEGF, NFκB, TNFα, PI3K, TGFβ, EGFR, Hypoxia, and MAPK in tumors compared to HCs (Fig. 2C). Among these, EGFR, Hypoxia, and MAPK signaling were particularly enhanced in tumor epithelial cells, while PI3K and TGFβ signaling was markedly activated in tumor fibroblasts. TNFα and NFκB pathways, central to inflammatory responses, showed notably elevated activity in tumor myeloid cells (Fig. 2C). Feature plots illustrated the expression patterns of representative upregulated DEGs across tumor and HC tissues (Fig. 2D). Together, these results provide comprehensive insight into cell type-specific transcriptional landscapes in SNSCC and highlight key molecular pathways potentially involved in its pathogenesis.

### Single-cell open chromatin landscapes reveal regulatory mechanisms associated with gene expression in SNSCC

To elucidate the regulatory epigenomic landscapes of SNSCC, we explored chromatin accessibility alterations across distinct cell types between SNSCC and HC tissues. Using stringent criteria (adjusted *p*-value < 0.05, > 2-fold differences in chromatin accessibility), we identified differentially accessible regions (DARs) within each cell type between tumor and HC tissues. Due to limited DARs (< 50 DARs) in immune cell populations, we focused our analysis on the three predominant cell types in SNSCC: epithelial cells, fibroblasts, and endothelial cells (Supplementary Fig. 3A). To understand the chromatin dynamics underlying transcriptional changes in SNSCC, we identified DARs linked to DEGs by comparing tumor cell types with their normal counterparts. Heatmap analysis revealed significant correlations between linked DARs and DEGs in each cell type (Fig. 3A), with approximately 10% of DARs closely associated with DEGs (Supplementary Fig. 3B). DEG-linked DARs were predominantly located in promoter-TSS, intron, and intergenic regions, suggesting that *cis*-regulatory elements such as promoters and enhancers play key roles in chromatin remodeling in SNSCC (Supplementary Fig. 3C). The remaining DARs mapped to TTS, 5’UTR, 3’UTR, exon, and non-coding regions. Functional analysis of cell type-specific accessible DARs associated with upregulated DEGs in tumors revealed transcriptional features consistent with our previous SNSCC characterization (Supplementary Fig. 3D). Within tumors, we observed significantly increased chromatin accessibility in DARs associated with cell type-specific genes. Tumor epithelial cells exhibited enhanced DNA accessibility in DARs linked to *DSG3*, *ENO1*, *KRT17*, and *NDRG1*. CAFs showed increased accessibility in DARs associated with *FAP*, *FBLN2*, *SFRP2*, and *TGFBI*, while tumor-infiltrating endothelial cells displayed elevated accessibility at DARs linked to *ANGPT2*, *COL4A1*, *PRDM1*, and *VMP1* (Fig. 3B). This enhanced chromatin accessibility strongly correlated with increased transcriptional activity of the corresponding genes (Supplementary Fig. 3E). Representative cell type-specific DARs and their adjacent gene expression patterns highlighted distinct functional roles within the TME (Fig. 3C). For example, Desmoglein-3 (DSG3) overexpression has been implicated in head and neck cancer progression^45^. FAP-expressing CAFs are known to promote tumor growth by fostering an immunosuppressive TME and contributing to immunotherapy resistance^46, 47, 48^. ANGPT2, a member of the angiopoietin family, has been reported to facilitate vascular destabilization and sprouting in tumors^49^. These results suggest that chromatin accessibility alterations drive the expression of genes crucial for SNSCC pathogenesis through cell type-specific regulatory mechanisms.

**Fig. 3.**
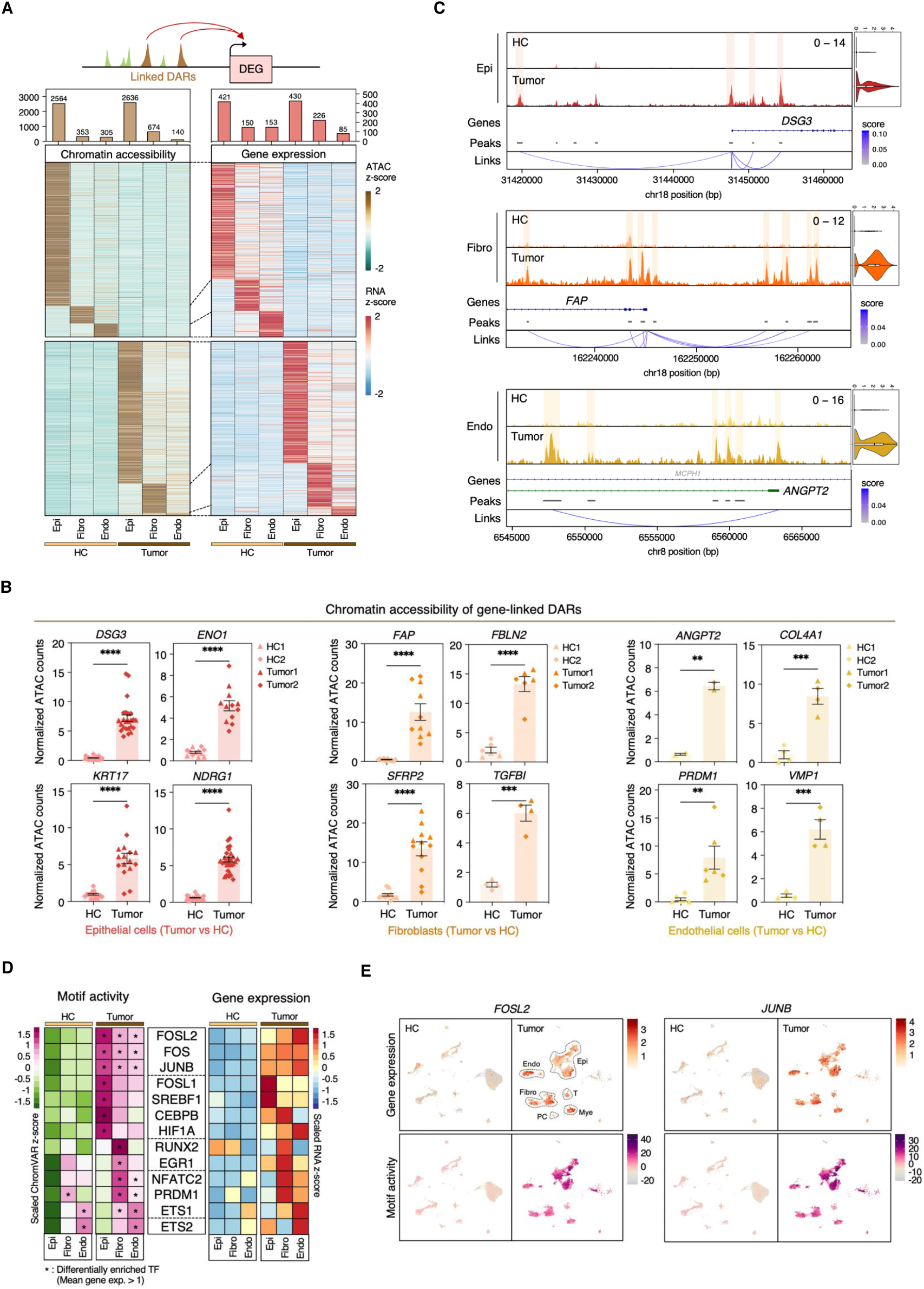
Single-nucleus chromatin accessibility profiling in SNSCC. **A** Schematic of linkage between differentially accessible regions (DARs) and DEGs (upper). Heatmap showing chromatin accessibility and gene expression of differential peaks linked with DEGs in epithelial cells, fibroblasts, and endothelial cells compared to their normal counterparts (lower). **B** Bar plots displaying the mean read counts of single-nucleus ATAC peaks for DARs linked to select DEGs in each cell type between SNSCC tumor and HC samples (Tumor1: CA5, Tumor2: CA8). Data are shown as mean ± SEM. *P* values were determined by two-tailed unpaired t-test. \**p* < 0.05, \*\**p* < 0.01, \*\*\**p* < 0.001, \*\*\*\**p* < 0.0001. **C** Coverage plots showing ATAC signals and gene expression levels of select DEGs identified in **B**, separated by cell types for SNSCC tumor (dark) and HC (light) samples, with peak coordinates and peak-gene links. **D** Heatmap showing chromVAR motif activity (left) and gene expression (right) of select differentially enriched TF binding motifs with upregulated gene expression in each cell type within SNSCC compared to their normal counterparts. **E** UMAP plots for gene expression (upper) and chromVAR motif activity (lower) of FOSL2 and JUNB, split by tissue type.

To investigate key transcription factors (TFs) in SNSCC, we identified differentially enriched motifs in each cell type between tumors and HCs (adjusted *p*-value < 0.05, > 0.58 log_2_-fold change) (Supplementary Fig. 3F). Each major cell type exhibited distinct TF motif activity patterns. For example, the lineage-specific TF PU.1 (SPI1) was strongly enriched in myeloid cells, while IRF4, involved in plasma cell differentiation, was enriched in plasma cells (Supplementary Fig. 3F). We focused on TFs with increased gene expression in tumors compared to normal tissues, identified among cell type-specific enriched motifs with moderate expression levels (mean expression > 1). The selected binding motifs, along with their ChromVAR motif accessibility and gene expression, are shown in Fig. 3D. We observed significant enrichment of the HIF1A motif in tumor epithelial cells, supporting the hypoxia-induced transcriptional responses identified in SNSCC. In tumor-associated fibroblasts, motifs such as RUNX2, EGR1, and NFATC2 were highly enriched, with RUNX2 known to regulate the functional specification of CAFs with tumorigenic potential^50^. ETS1 is considered an important TF in endothelial cells, associated with angiogenesis^51, 52, 53^. ETS family TFs, particularly ETS1 and ETS2, showed strong gene expression and motif enrichment in tumor endothelial cells. Notably, AP-1 motifs, including FOS, FOSL2, and JUNB, were prominently enriched in tumor epithelial cells and broadly enriched across all tumor cell types (Fig. 3D). The expression levels of *FOSL2* and *JUNB* were upregulated in tumors compared to normal tissues (Supplementary Fig. 3G). Furthermore, the correlation between enhanced gene expression of *FOSL2* and *JUNB* in tumor cell compartments and their motif activity suggests that AP-1 plays a crucial role in SNSCC regulatory mechanisms (Fig. 3E). Together, these results demonstrate that integrating chromatin accessibility profiling with motif analysis reveals cell type-specific regulatory landscapes, highlighting their significant contributions to the complex transcriptional programs driving SNSCC pathogenesis.

### SNSCC exhibits globally altered DNA methylome profiles

DNA methylation has been shown to serve as a potent epigenetic mechanism for sinonasal cancer classification^22^, and our bulk RNA-seq analysis revealed enriched DNA methylation-related ontologies among upregulated genes in SNSCC. To examine global DNA methylation changes in SNSCC, we performed methylation profiling array (EPIC array) analyses on tumor (*n* = 3) and normal (*n* = 3) samples from SNSCC patients, which were included in our bulk RNA-seq analysis. We integrated these data with public EPIC array datasets comprising 35 SNSCC and 8 normal sinonasal tissues from non-cancer patients (GSE189778). Multi-dimensional scaling (MDS) analysis of the 1000 most variable CpG sites effectively distinguished normal and tumor groups. Our DNA methylation array data clearly separated normal and tumor samples and showed patterns similar to public datasets, validating the reliability of this integrative analysis (Supplementary Fig. 4A).

Differential methylation analysis identified 39,526 hypomethylated and 14,605 hypermethylated CpG sites in tumors compared to normal tissues (adjusted *p*-value < 0.05, > 20% β-value differences) (Fig. 4A and Supplementary Fig. 4B). Examination of differentially methylated positions (DMPs) across genomic regions revealed that hypomethylated CpGs were enriched in intergenic regions (IGRs), while hypermethylated CpGs were more frequent near transcription start sites (TSSs), including TSS200, TSS1500, 5’UTR, and 1st exon (Fig. 4B). Additionally, hypomethylated CpGs were highly enriched in opensea regions, whereas hypermethylated CpGs were predominantly located within CpG islands (Fig. 4C).

**Fig. 4.**
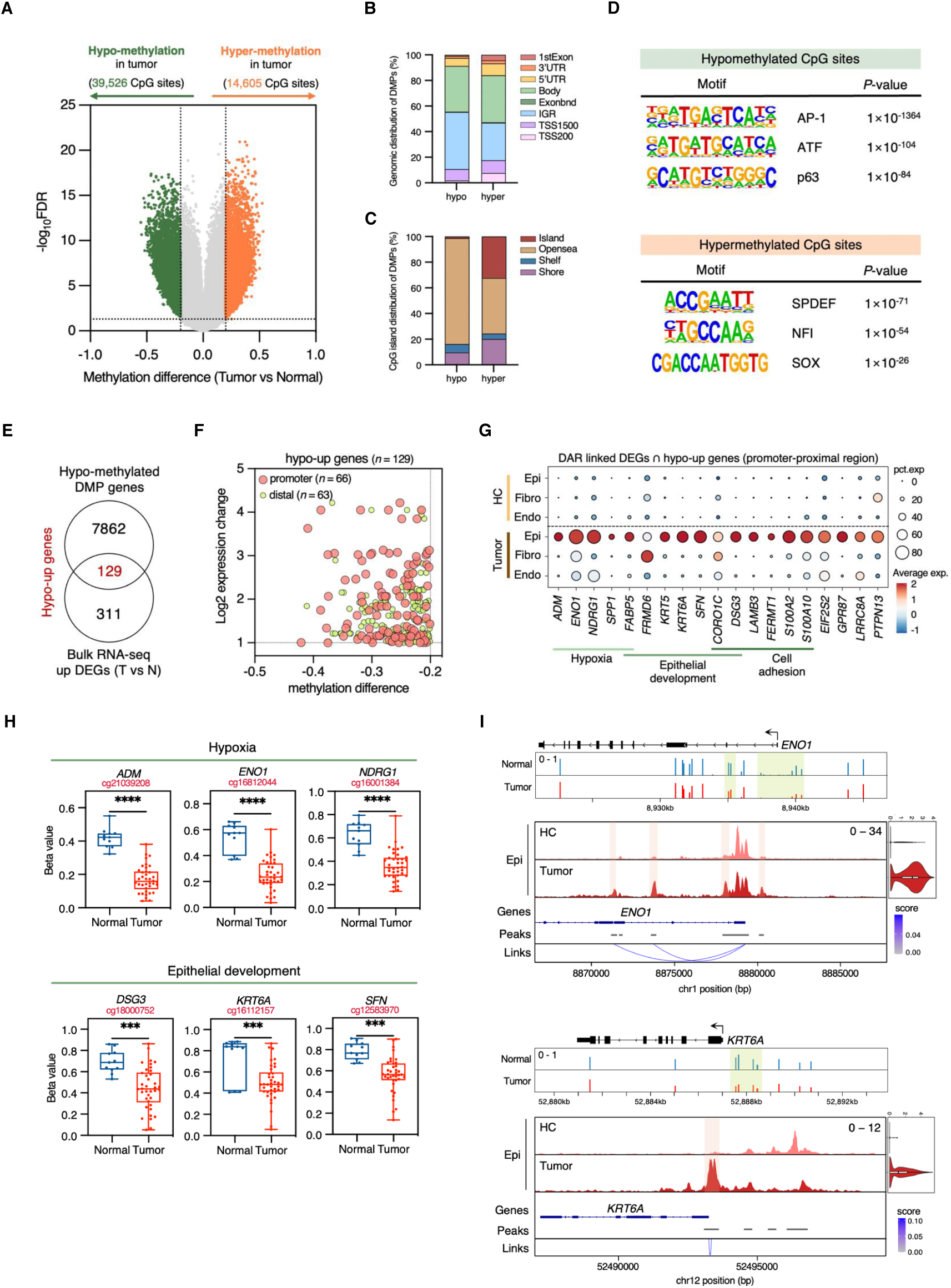
Extensive epigenetic reprogramming with gene expression changes in SNSCC. **A** Volcano plot illustrating the differentially methylated positions (DMPs) between SNSCC tumors and normal nasal tissues. Significant DMPs were defined by a β-value threshold greater than 0.2 and an adjusted *p*-value less than 0.05. Gray dots represent non-significant CpG sites. **B** Genomic distribution of hypo- and hypermethylated DMPs. **C** Distribution of CpG islands (CGIs) of hypo- and hypermethylated DMPs. **D** Significantly enriched binding motifs in hypo- (upper) and hypermethylated (lower) DMPs revealed by de novo motif analysis using HOMER software. **E** Venn diagram showing the number of hypomethylated upregulated (hypo-up) genes in SNSCC tumor compared to normal samples. **F** Scatter plot displaying the mean difference in β-value and log2 fold change in gene expression of 129 hypo-up genes, with genomic regions including promoter (red) and distal (green). Each dot represents a gene-CpG pair. **G** Gene expression levels of DAR-linked DEGs in tumor-specific cells identified in Fig. 3A intersecting with 66 hypo-up genes in promoter-proximal regions in each cell type within SNSCC tumor and HC samples. Functional features related to intersecting genes are presented at the bottom. **H** Box plots showing β-values of DMPs of select hypo-up genes identified in **G** in SNSCC tumor and normal samples. Each dot represents one donor. Boxes encompass the 25th to 75th percentiles, whiskers extend from min to max, and the center line indicates the median. *P* values were determined by two-tailed unpaired t-test. \**p* < 0.05, \*\**p* < 0.01, \*\*\**p* < 0.001, \*\*\*\**p* < 0.0001. **I** Genome browser track of DNA methylation levels at *ENO1* and *KRT6A* loci in SNSCC tumor (red) and normal (blue) samples (upper). Coverage plots showing ATAC signals and gene expression levels of *ENO1* and *KRT6A* in epithelial cells within SNSCC tumor (dark) and HC (light) samples with peak coordinates and peak-gene links (lower).

Since DNA methylation states can regulate gene transcription by modulating TF binding affinity, we investigated enriched TF binding motifs associated with DMPs in SNSCC using HOMER. Motif analysis revealed significant enrichment of AP-1, KLF, and p63 motifs in hypomethylated CpG sites (Fig. 4D). Further analysis showed enrichment of AP-1 and KLF motifs in hypomethylated CpGs within promoter-proximal regions (TSS, 5’UTR, and 1st exon), suggesting these motifs may substantially influence transcriptional regulation (Supplementary Fig. 4C). In contrast, hypermethylated DMPs showed enrichment for SPDEF, NFI, and SOX motifs, accompanied by decreased expression of NFI family members, including *NFIB* and *NFIX*, which are essential for normal development and possess tumor-suppressive potential in diverse cancers (Fig. 4D and Supplementary Fig. 4D)^54^. Notably, these observations aligned with our previous TF enrichment analysis from snRNA/ATAC-seq data (Fig. 3E and Supplementary Fig. 4E). These findings suggest that TFs such as FOSL2 and JUNB may serve as crucial mediators in the broader epigenetic changes driving SNSCC tumorigenesis.

### Epigenetic regulation drives transcriptional changes in SNSCC

Considering the well-established inverse relationship between DNA methylation and gene expression, we integrated DMP-associated genes and bulk DEGs in SNSCC to identify two categories of genes: 129 hypomethylated-upregulated (hypo-up) and 102 hypermethylated-downregulated (hyper-down) genes, each containing one or more DMPs (Fig. 4E and Supplementary Fig. 4F). To investigate the impact of aberrant DNA methylation on gene activation, we focused on hypo-up genes. Among 264 differentially methylated CpG sites corresponding to the 129 hypo-up genes, we observed 110 hypomethylated CpG sites annotated on 66 genes in promoter-proximal regions and 154 CpG sites annotated on 63 genes in distal regions outside the proximal promoter (Fig. 4F). Given the strong correlation between promoter DNA methylation and transcriptional activity, we explored potential TF motifs enriched in the hypomethylated DMPs within the promoter-proximal regions of hypo-up genes. We identified AP-1 and p53 family binding motifs enriched in hypomethylated CpG sites corresponding to upregulated genes in SNSCC (Supplementary Fig. 4G).

Since reduced DNA methylation is associated with increased chromatin accessibility and facilitates transcriptional activation, we intersected the 66 hypo-up genes showing promoter hypomethylation with DEGs linked to DARs in tumors versus HCs to discover hypomethylated genes regulated by accessible chromatin regions. We found that most of the 19 overlapping genes exhibited higher expression levels in tumor epithelial cells (Fig. 4G). Specifically, significant reductions in DNA methylation levels were observed at CpG sites within the promoters of genes involved in hypoxia (*ADM*, *ENO1*, and *NDRG1*) and epithelial development (*DSG3*, *KRT6A*, and *SFN*) in SNSCC (Fig. 4H). For example, elevated expression levels of *ENO1* and *KRT6A* within tumor epithelium were attributed to increased chromatin accessibility and loss of DNA methylation in their promoter regions (Fig. 4I). These findings collectively highlight the impact of epigenomic reprogramming on the transcriptional landscape of SNSCC, particularly in tumor epithelium.

### Molecular profiling identifies clinically relevant malignant epithelial subpopulations in SNSCC

To further elucidate the dynamics of epithelial cells in tumor and HC tissues, we reclustered the epithelial compartment into 15 clusters (Supplementary Fig. 5A, B). We annotated 12 cell types based on marker gene expression, including basal cells (*KRT15* and *TP63*), ciliated cells (*FOXJ1* and *PIFO*), goblet cells (*MUC5AC* and *SCGB1A1*), myoepithelial cells (MECs; *ACTA2* and *MYH11*), mucous cells (*MUC5B* and *BPIFB2*), serous cells (*LYZ* and *LTF*), tuft cells (*POU2F3* and *TRPM5*), and five tumor cell subpopulations (TC1-5) predominantly composed of cells from tumor tissues (Fig. 5A, B and Supplementary Fig. 5C, D). Most epithelial compartments, except for TC subpopulations and tuft cells, were found in HC tissues, and the TC3 cluster included some epithelial cells from control nasal mucosa (Fig. 5B and Supplementary Fig. 5D). TC subpopulations TC1-4 were present in both tumor samples, while TC5 was predominantly detected in one patient sample (Fig. 5B). Genes significantly upregulated in head and neck squamous cell carcinoma (HNSC) tumors, such as *KRT6A* and *SFN*, were highly expressed across all tumor cell subpopulations. In contrast, *AZGP1* and *PROM1*, significantly downregulated in HNSC patients, showed notable expression in normal nasal epithelial compartments, such as mucous and serous cells (Fig. 5C, D). Moreover, scores of the top 100 tumor-upregulated DEGs from bulk RNA-seq were significantly elevated in all TC clusters, particularly TC1 and TC2, whereas scores of the top 100 downregulated DEGs were increased in control tissue-enriched epithelial populations (Supplementary Fig. 5E). Inferred copy number variations (CNVs) analysis using ciliated, mucous, and serous cells as references clearly separated TC clusters from other populations (Fig. 5E). TC clusters displayed higher CNVs compared to other epithelial cell subtypes, with TC1, TC2, and TC4 showing the highest levels (Fig. 5F). These findings confirmed that TC subclusters represent malignant cells within SNSCC.

**Fig. 5.**
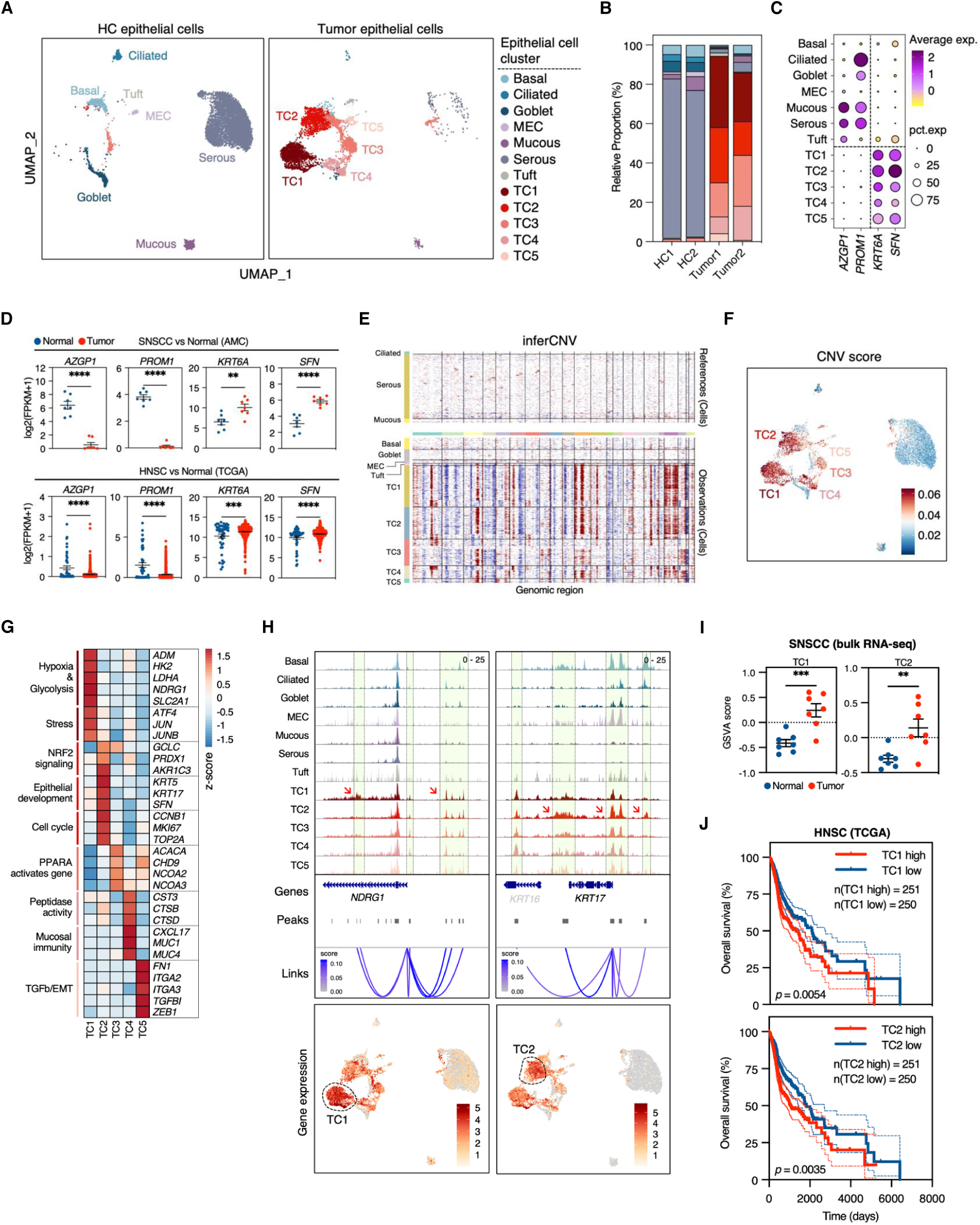
Heterogenous tumor epithelial subpopulations show distinct characteristics. **A** UMAP plots of epithelial compartments in SNSCC tumor and HC samples computed by WNN, colored by cell type. **B** Proportion of epithelial cell states in each sample (Tumor1: CA5, Tumor2: CA8). **C** Dot plot showing gene expression levels of select genes that are down- (*AZGP1* and *PROM1*) or upregulated (*KRT6A* and *SFN*) DEGs from bulk RNA-seq analysis in each epithelial cell subpopulation. **D** Expression levels of *AZGP1*, *PROM1*, *KRT6A*, *SFN* in SNSCC tumor versus normal samples from bulk RNA-seq (upper) and head and neck squamous cell carcinoma (HNSC) versus normal samples from The Cancer Genome Atlas (TCGA) dataset (lower). Each dot represents one donor. Data are shown as mean ± SEM. *P* values were determined by two-tailed unpaired t-test. \**p* < 0.05, \*\**p* < 0.01, \*\*\**p* < 0.001, \*\*\*\**p* < 0.0001. **E** Inferred copy number variations (CNVs) of all malignant cell clusters using ciliated, mucous, and serous cells enriched in normal mucosa as reference cells, analyzed by inferCNV. **F** UMAP plot of epithelial populations colored by inferred CNV score. **G** Heatmap showing the expression levels and functional characteristics of select genes identified in DEGs across malignant cell subclusters (TC1-5). **H** Coverage plots showing ATAC signals of *NDRG1* and *KRT17* in epithelium with peak coordinates and peak-gene links colored by epithelial cell subcluster (upper). UMAP plots displaying gene expression levels of *NDRG1* and *KRT17* (lower). **I** Signature scores of TC1 and TC2 in SNSCC tumor versus normal samples from bulk RNA-seq. Each dot represents one donor. Data are shown as mean ± SEM. *P* values were determined by two-tailed unpaired t-test. \**p* < 0.05, \*\**p* < 0.01, \*\*\**p* < 0.001, \*\*\*\**p* < 0.0001. **J** Kaplan-Meier curves of overall survival according to high and low groups in HNSC patients from TCGA dataset stratified by TC1 (upper) and TC2 (lower) signature scores (50th percentile). *P* values were determined by log-rank test.

To dissect intratumoral heterogeneity in SNSCC, we compared gene expression profiles across TC clusters. GO analysis using DEGs (adjusted *p*-value < 0.05, > 0.58 log_2_-fold change) revealed functional differences among the five TC subpopulations (Supplementary Fig. 5F, G). TC1 was enriched in pathways associated with hypoxia, glycolysis, and stress response (*ATF4*, *JUN*, and *JUNB*), indicating a core tumor-promoting role in metabolic stress. TC2 was characterized by expression of genes related to epithelial development (*KRT5*, *KRT17*, and *SFN*), cell cycle (*CCNB1*, *MKI67*, and *TOP2A*), and NRF2 pathway components such as peroxiredoxin (*PRDX1*) and reductases (*AKR1C3*). TC3 showed upregulation of PPARA-regulated genes (*NCOA2* and *NCOA3*) and DNA damage response genes. TC4 featured genes involved in peptidase activity (*CTS3*, *CTSB*, and *CTSD*) and mucosal immunity, including mucosal defense (*MUC1* and *MUC4*) and mucosal chemokine (*CXCL17*). TC5 exhibited molecular features related to epithelial-mesenchymal transition (EMT), including extracellular matrix (ECM)-receptor interaction, TGF-β signaling, and wound healing, supported by expression of EMT markers (*FN1* and *ZEB1*) and integrins (*ITGA2* and *ITGA3*) (Fig. 5G and Supplementary Fig. 5G). TC populations displayed distinct chromatin accessibility states compared to normal-enriched cells (Fig. 5H). For example, TC-specific peaks with increased accessibility were observed in *NDRG1*-related *cis*-regulatory elements across all TC subclusters, with some peaks showing higher accessibility in TC1 (red arrows), correlating with gene expression patterns. Similarly, we identified highly accessible peaks at the *KRT17* locus in TC populations, with certain peaks showing higher accessibility in TC2 (red arrows) (Fig. 5H).

Intersection of TC subcluster DEGs with upregulated genes from bulk RNA-seq revealed that TC1 and TC2 showed particularly abundant overlap with SNSCC-overexpressed genes (Supplementary Fig. 5H). Additionally, TC1 and TC2 signature scores were most significantly enhanced in SNSCC tumor tissues (Fig. 5I and Supplementary Fig. 5I). These findings demonstrate both the widespread presence of these malignant subsets across SNSCC patients and their potential role in tumor development. Notably, TC1-high and TC2-high groups exhibited poorer clinical outcomes compared to their respective low-expression groups in The Cancer Genome Atlas (TCGA) HNSC data, while no significant differences in overall survival rates were observed among other TC clusters (Fig. 5J and Supplementary Fig. 5J). Together, these findings provide insight into tumor cell heterogeneity in SNSCC, with TC1 and TC2 malignant population signatures emerging as critical indicators of clinical outcomes.

### Hypoxic tumor cells promote angiogenesis through paracrine signaling in SNSCC

To determine crucial paracrine signaling by malignant cells in the SNSCC TME, we investigated cell-cell interactions between major cell types in HC and tumor tissues. We observed prominent interactions between epithelial and endothelial cells in tumors, but sparse connections in HC tissues, as indicated by edge thickness (Fig. 6A). Notably, our bulk RNA-seq data showed significantly upregulated Hypoxia and VEGF signaling pathways in tumor compared with normal tissues (Supplementary Fig. 6A). Given that hypoxic microenvironments promote aggressive tumor phenotypes and impede therapeutic responses through abnormal vascular development^34, 55^, we focused on secreted factors from TC1 population representing hypoxic tumor cells, which showed activated hypoxia and glycolysis pathway signatures (Supplementary Fig. 6B).

**Fig. 6.**
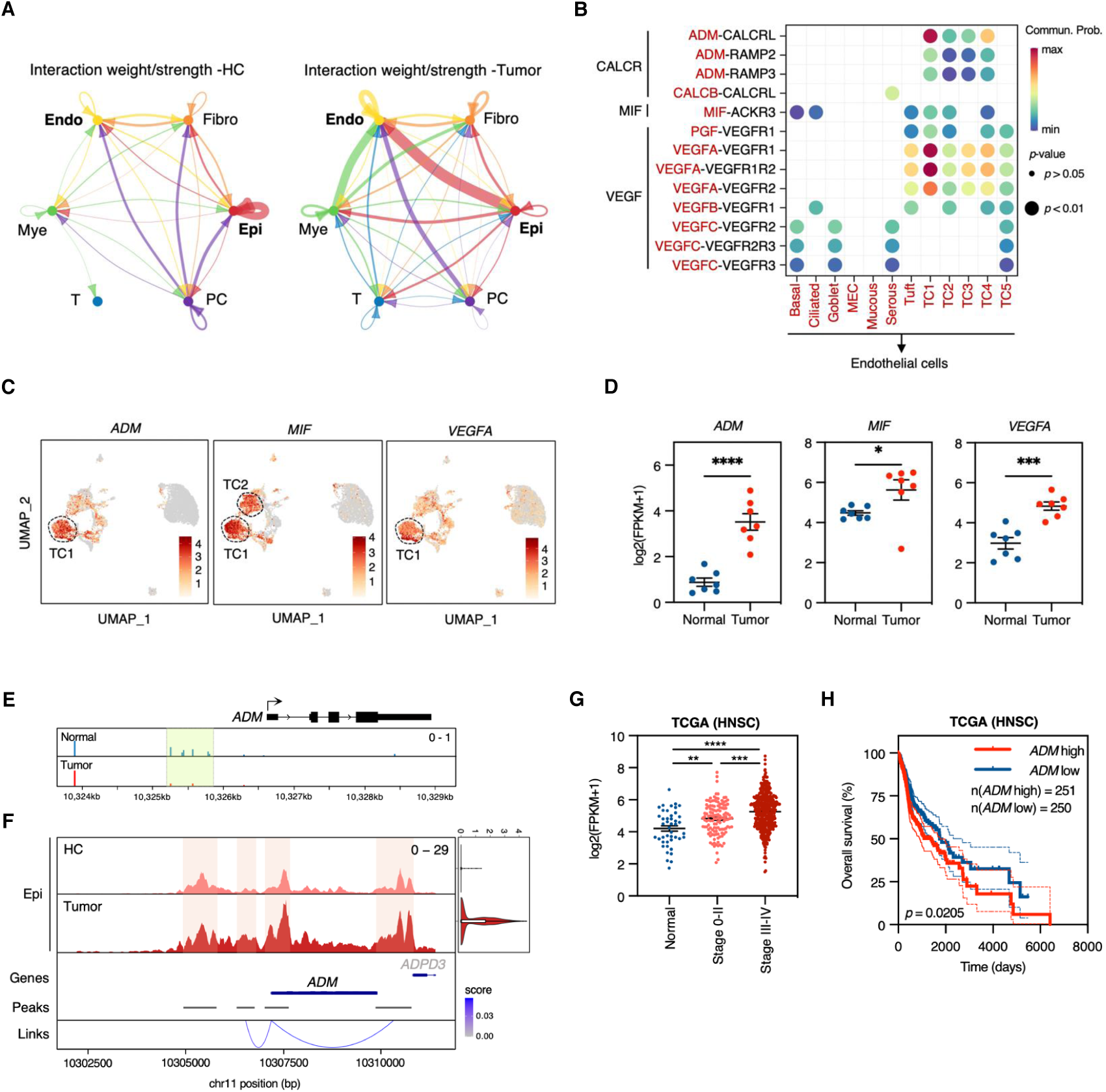
Hypoxic tumor cells promote angiogenesis by secreting angiogenic factors. **A** Circle plots representing cell-cell communication networks for major cell types in SNSCC tumor and HC samples using CellChat. Interactions include secreted signaling from the CellChat database. Interaction strength indicated by the connecting edge thickness. **B** Bubble plot of communication probability for significant ligand-receptor pairs associated with CALCR, MIF, and VEGF signaling pathways from all epithelial cell subclusters (source) to endothelial cells (target). **C** UMAP plots showing gene expression levels of *ADM*, *MIF*, and *VEGFA* in epithelium compartments. **D** Expression levels of *ADM*, *MIF*, and *VEGFA* in SNSCC tumor versus normal samples from bulk RNA-seq. Each dot represents one donor. Data are shown as mean ± SEM. *P* values were determined by two-tailed unpaired t-test. \**p* < 0.05, \*\**p* < 0.01, \*\*\**p* < 0.001, \*\*\*\**p* < 0.0001. **E** Genome browser track of DNA methylation levels at the *ADM* locus in SNSCC tumor (red) and normal (blue) samples. **F** Coverage plots showing ATAC signals and gene expression levels of *ADM* in epithelial cells within SNSCC tumor (dark) and HC (light) samples with peak coordinates and peak-gene links. **G** Expression levels of *ADM* in HNSC samples from TCGA dataset grouped by normal, stage 0-II, and stage III-IV. Each dot represents one donor. Data are shown as mean ± SEM. *P* values were determined by two-way ANOVA. \**p* < 0.05, \*\**p* < 0.01, \*\*\**p* < 0.001, \*\*\*\**p* < 0.0001. **H** Kaplan-Meier curves of overall survival according to high and low groups in HNSC patients from TCGA dataset, stratified by *ADM* expression levels (50th percentile). *P* values were determined by log-rank test.

Cell-cell interaction analysis between epithelial cells (source) and endothelial cells (target) revealed enrichment of CALCR, MIF, and VEGF signaling pathways in tumors, with strong interactions originating from TC1 populations. In contrast, ncWNT signaling was enriched in control nasal tissues, primarily sourced from plasma cells (Supplementary Fig. 6C, D). MIF signaling was strongly activated between both TC1 and TC2 clusters and myeloid cells, suggesting that tumor-derived MIF could influence pro-tumor myeloid activation (Supplementary Fig. 6D). Analysis of significant ligand-receptor pairs revealed that the TC1 cluster predominantly interacts with endothelial cell receptors via ADM-CALCRL, ADM-RAMP2, ADM-RAMP3, MIF-ACKR3, VEGFA-VEGFR1, and VEGFA-VEGFR2 pairs (Fig. 6B).

Consistently, expression of pro-angiogenic factors *ADM*, *MIF*, and *VEGFA* was markedly enhanced in the TC1 cluster (Fig. 6C) and significantly upregulated in tumor tissues compared with normal tissues in our bulk transcriptome data (Fig. 6D).

Adrenomedullin (ADM), induced under hypoxic conditions, can promote angiogenesis, leading to cancer progression and carcinogenesis^56^. We found that DNA methylation levels at *ADM* loci were considerably decreased in SNSCC tumor compared with normal control tissues (Fig. 6E). Moreover, chromatin accessibility around the *ADM* gene locus was increased in tumor epithelial cells compared to HC tissues (Fig. 6F), suggesting that *ADM* expression is regulated by epigenetic changes in SNSCC. Using the TCGA-HNSC cohort, we validated the relationship between *ADM* expression and clinicopathological features. *ADM* expression significantly increased with clinical stage in HNSC patients (Fig. 6G) and strongly correlated with TC1 signature genes, such as *NDRG1*, and *VEGFA* expression (Supplementary Fig. 6E). Furthermore, patients with high *ADM* expression showed significantly shorter overall survival than those with low expression (Fig. 6H). These findings indicate that endothelial cells respond to angiogenic stimuli (ADM, MIF, and VEGFA) secreted by hypoxia-related tumor cell populations, thereby promoting angiogenesis and potentially contributing to SNSCC pathogenesis. These results also suggest that *ADM* could serve as a promising therapeutic target in SNSCC.

### Endothelial tip cells stimulated by hypoxic tumor cells promote SNSCC progression

To elucidate endothelial cell diversity and angiogenic potential, we identified six distinct endothelial cell clusters, excluding doublets, within SNSCC tumor and HC tissues (Fig. 7A and Supplementary Fig. 7A, B). These clusters were annotated into five distinct endothelial subtypes based on marker gene expression: tip (*CXCR4*, *DLL4*, and *ESM1*), immature (*ENG*, *HSPG2*, and *VWA1*), venous (*CLU* and *VWF*), lymphatic (*LYVE1* and *PROX1*), and arterial (*CXCL12* and *GJA4*) endothelial cells. Immature cells also expressed stalk-like marker genes such as *ACKR1*, *SELE*, and *SELP* ^57^. Venous cells comprised two subsets with similar expression patterns (Fig. 7B). The EC1 (tip) and EC2 (immature) clusters were exclusively enriched in tumor tissues, while venous cells were enriched in HC tissues (Fig. 7C), consistent with previous reports on endothelial cell phenotypes in normal adjacent and tumor samples from lung, breast, and gastric cancer patients^58, 59, 60^.

**Fig. 7.**
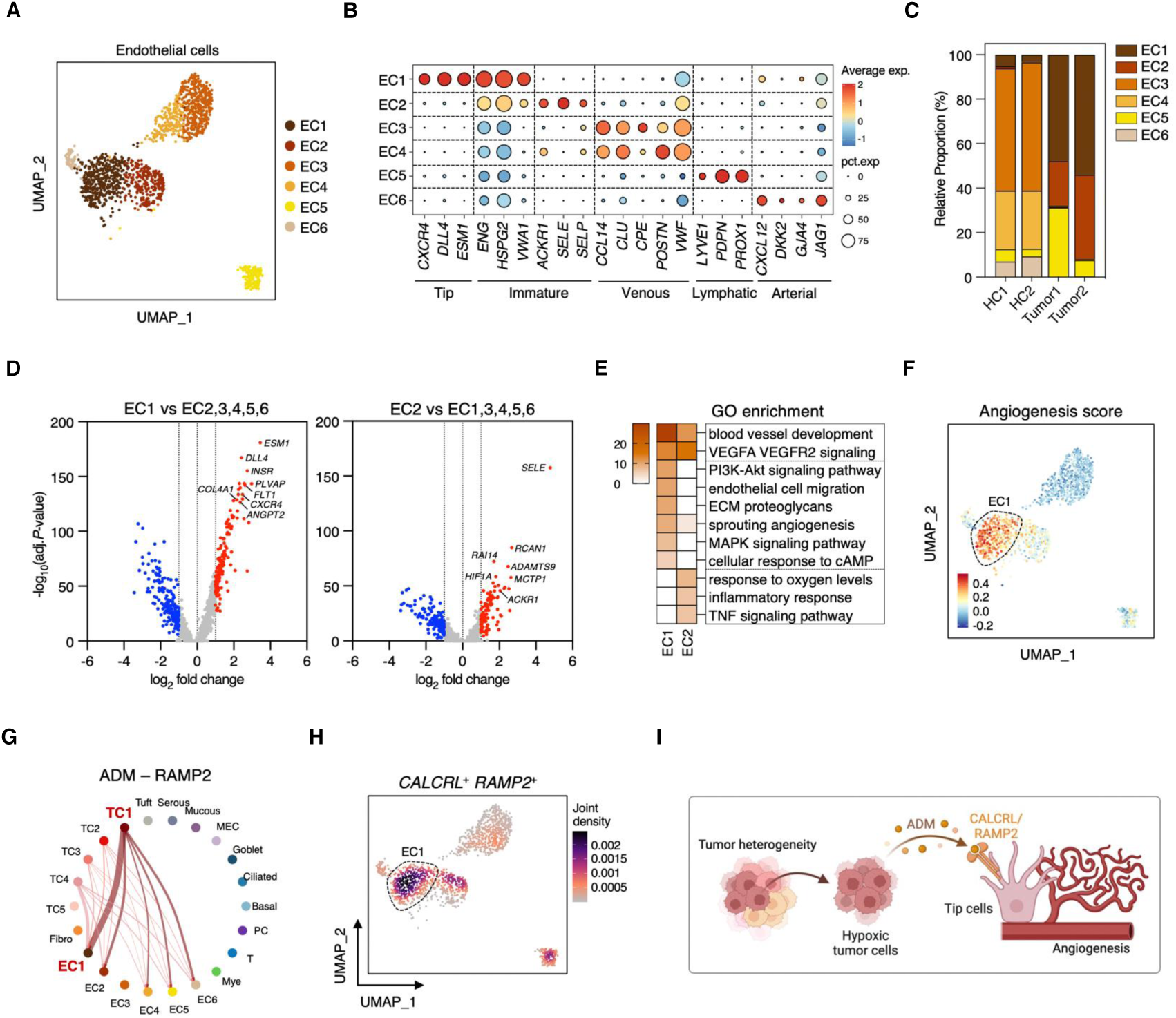
Endothelial tip cell activation by hypoxic tumor cells induces progression of SNSCC. **A** UMAP visualization of endothelial compartments in SNSCC tumor and HC samples by WNN analysis, colored by subclustering. **B** Dot plot of expression levels of marker genes for each endothelial cell subpopulation. **C** Proportion of endothelial cell states in each sample (Tumor1: CA5, Tumor2: CA8). **D** Volcano plots of the transcriptional changes in EC1 (left) and EC2 (right) compared to other endothelial subclusters. Significant DEGs were determined if log2 fold change is greater than 1 and adjusted *p*-value is less than 0.05. Gray dots represent non-significant genes. **E** Heatmap representing the *p*-value significance of enriched GO terms for upregulated genes in EC1 and EC2 cluster identified in **D**. **F** UMAP plot of angiogenesis signature scores in endothelial cells. **G** Circos plot of inferred ADM-RAMP2 interactions between all epithelial and endothelial states. The thickness of edge representing the communication probability. **H** Density plot of *CALCRL*^+^ *RAMP2*^+^ cells in endothelial compartments. **I** Schematic representation of the proposed mechanism whereby hypoxic tumor cells secrete ADM, which binds to receptors (CALCRL/RAMP2) on tumor-specific tip cells to promote angiogenic sprouting. Created with Biorender.com.

To identify transcriptional changes in tumor-enriched endothelial cells, we examined DEGs between EC1 and other endothelial subpopulations (EC2-6), and similarly compared EC2 against the remaining subpopulations (Fig. 7D). GO analysis revealed angiogenic features (e.g., blood vessel development and VEGFA VEGFR2 signaling) in tumor-associated endothelial compartments (Fig. 7E). EC1 showed strong enrichment in angiogenesis-related pathways (e.g., PI3K-Akt and MAPK) and ECM proteoglycan, with high expression of genes involved in angiogenic regulation (*ANGPT2* and *PLVAP*), ECM (*COL4A1* and *COL4A2*), and integrins (*ITGA1* and *ITGA8*) (Fig. 7D, E and Supplementary Fig. 7C). These findings indicate that EC1 possesses the highest angiogenic potential. In contrast, EC2 was associated with oxygen response (*HIF1A* and *NDRG1*) and immune activation, including immune cell recruitment (*ACKR1* and *SELP*)^61, 62^ and inflammation (*NFKBIA* and *ICAM1*) (Fig. 7D, E and Supplementary Fig. 7C). Consistently, EC1 exhibited higher angiogenesis signature scores (Fig. 7F).

To investigate which endothelial subset interacts most strongly with hypoxic tumor cells, we focused on the CALCRL/RAMP2 complex^63^, the primary receptor for ADM functionality which is involved in tumorigenic pathways including cell proliferation and angiogenesis^64^. While all endothelial subclusters showed robust ADM-CALCRL interactions with TC1 (Supplementary Fig. 7D), strong ADM-RAMP2 interactions were specifically observed between TC1 and EC1 clusters (Fig. 7G). Moreover, *CALCRL* and *RAMP2* double-positive cells were predominantly enriched in EC1 (Fig. 7H). Gene expression levels of *CALCRL* significantly correlated with *RAMP2* expression in the HNSC patient cohort (Supplementary Fig. 7E). We also identified strong MIF-ACKR3, MIF-(CD73+CXCR4), VEGFA-VEGFR1, and VEGFA-VEGFR2 ligand-receptor interactions between TC1 and EC1 (Supplementary Fig. 7F). Consistently, expression levels of *ACKR3*, *CXCR4*, *FLT1* (VEGFR1), and *KDR* (VEGFR2) were notably enhanced in EC1 (Supplementary Fig. 7G). These findings suggest that secreted factors such as ADM from hypoxic tumor cells interact with receptors on tumor-specific tip cells, stimulating angiogenic sprouting and promoting SNSCC progression (Fig. 7I).

## Discussion

The molecular underpinnings of SNSCC have remained poorly understood due to its rarity and limited availability of patient samples for comprehensive analysis. Our integrative multi-omic profiling reveals comprehensive molecular characteristics and cell type-specific signatures of the SNSCC landscape. Through integration of bulk and single-cell approaches, we identified core oncogenic programs, epigenetic regulation of transcriptional changes, and heterogeneous cellular components driving SNSCC pathogenesis.

A central finding of our study is the identification of distinct malignant cell populations with clinical relevance. The hypoxic (TC1) and proliferative (TC2) clusters not only define the cellular heterogeneity within SNSCC but also highlight how different tumor cell states may contribute to disease progression. The strong association between these populations and poor clinical outcomes in the HNSC dataset, suggests they may represent key drivers of aggressive disease behavior. Particularly noteworthy is the TC1 cluster’s hypoxic signature, which builds on previous observations of oxygen-deprived regions in head and neck cancers^36, 65^, but provides detailed molecular characterization of hypoxia-adapted tumor cells. These hypoxic features are prominently represented among tumor-upregulated genes compared to normal mucosa in our bulk transcriptomic data. The robust glycolytic programming in TC1 cells likely represents an adaptation to sustained hypoxic stress, enabling these cells to survive in nutrient-poor conditions while maintaining aggressive phenotypes. This finding has important therapeutic implications, as hypoxic tumor cells are often resistant to conventional treatments. The co-existence of TC1 and TC2 populations suggests that therapeutic strategies targeting either population alone may be insufficient, supporting the development of combination approaches that address both hypoxic and proliferative tumor compartments.

Our discovery of the TC1-EC1 signaling axis reveals a previously unrecognized mechanism driving tumor angiogenesis in SNSCC. Endothelial tip cells are enriched not only in SNSCC but also in other malignancies^58, 59, 66^. Integrative analyses across multiple cancer types have demonstrated that enrichment of endothelial tip cells correlates with poor clinical outcomes, with malignant cells and myeloid populations serving as primary sources of pro-angiogenic factors^66^. The preferential interaction between hypoxic tumor cells (TC1) and endothelial tip cells (EC1), mediated through hypoxia-induced factors including ADM, MIF, and VEGFA, represents a key mechanism for tumor-driven angiogenesis in SNSCC. The identification of ADM as a central mediator in this axis is particularly significant, suggesting opportunities for selective disruption of tumor vasculature. ADM expression strongly correlates with clinical outcomes in HNSC patients, indicating its particular relevance in aggressive disease. The epigenetic regulation of ADM further suggests potential therapeutic strategies through epigenetic modulation. Recent evidence demonstrates that ADM secreted by hypoxic TAMs destabilizes endothelial cell adherens junctions, indicating that ADM blockade could normalize vascular integrity and enhance drug delivery^67^. This hypoxia-angiogenesis axis presents a compelling therapeutic opportunity, particularly through targeting ADM or its receptors in SNSCC patients with pronounced hypoxic characteristics.

Our epigenomic analysis reveals a complex regulatory landscape in SNSCC that extends beyond previously reported DNA methylation patterns^22^. Through integrated analysis of chromatin accessibility and DNA methylation states, we identify cell type-specific regulatory programs that distinguish tumor-associated populations from their normal counterparts. We identified cell type-specific open chromatin landscapes in epithelial cells, fibroblasts, and endothelial cells within the TME that differ substantially from their normal counterparts. The consistent enrichment of AP-1 motifs in both hypomethylated regions and accessible chromatin across multiple cell types suggests AP-1 transcription factors serve as master regulators of the SNSCC cellular phenotype. This finding has immediate therapeutic relevance given recent advances in developing small molecule inhibitors targeting AP-1 family members^68, 69^.

The immune landscape of SNSCC is characterized by distinctive features that may influence treatment response. We observed substantial infiltration of myeloid cells expressing tumor-promoting pathways within the TME. These findings align with previous observations of increased myeloid cell proportions and diminished T cell presence in recurrent SNSCC^70^, suggesting that TME composition may critically influence disease progression and clinical outcomes. While the depth of immune cell characterization in our single-nucleus analysis was constrained by cell numbers, the identified patterns suggest that successful immunotherapeutic strategies may require specific targeting of the myeloid compartment. Future analyses of flow-sorted CD45^+^ cell populations will be essential to fully map the immune cell dynamics and identify optimal intervention points. Understanding the molecular mechanisms driving this unique immune landscape, particularly the interplay between myeloid cells and other TME components, could reveal novel therapeutic opportunities in SNSCC.

These findings have several important implications for therapeutic development. First, the identification of distinct tumor subpopulations suggests that combination therapies targeting multiple cellular states may be necessary for effective treatment. Second, the characterization of the TC1-EC1 signaling axis provides a rational basis for developing targeted anti-angiogenic approaches, potentially through ADM pathway inhibition. Third, the epigenetic landscapes we identified suggest opportunities for epigenetic therapy, particularly through modulation of AP-1-dependent transcription.

In conclusion, our integrated multi-omic analysis of SNSCC reveals the complex molecular architecture driving tumor progression. Through parallel analysis of transcriptional programs, epigenetic regulation, and cellular interactions, we provide a high-resolution map of the SNSCC tumor ecosystem. Key discoveries include the identification of clinically relevant tumor subpopulations (TC1 and TC2), characterization of a critical hypoxia-driven angiogenesis axis mediated by TC1-EC1 interactions, and elucidation of epigenetic mechanisms coordinating cell state-specific gene expression. The strong correlation between TC1/TC2 signatures and clinical outcomes suggests these populations could serve as prognostic biomarkers, while the ADM-dependent angiogenic axis presents an immediately targetable pathway. Our findings suggest several therapeutic strategies, including combination approaches targeting both hypoxic and proliferative tumor compartments, selective disruption of tumor angiogenesis through ADM pathway inhibition, and modulation of epigenetic states to normalize cell function. Future studies should focus on functional validation of these pathways, development of targeted therapeutic approaches.

## Methods

### Patient sample collection

Tumor samples were obtained from Korean adult patients with SNSCC at the time of surgical resection. Briefly, tumors were resected, flash frozen, and stored at a temperature of less than -80°C until RNA and DNA were extraction. Each frozen tumor specimen had an accompanying normal tissue from the contralateral nasal cavity. The weight of frozen tumor and normal tissue was at least 150 mg each, and usually less than 200 mg. This study was conducted in accordance with the Declaration of Helsinki and approved by the Institutional Review Board of Asan Medical Center (2019-0708). All participants provided written informed consent for inclusion before they participated in the study. Similar to clinical populations all patients were treated with curative intent. All cases were collected regardless of surgical stage or histologic grade and staged according to the 8th edition of the American Joint Committee on Cancer (AJCC).

Complete clinical data elements were collected including gender, age at diagnosis, tumor subsite, year of tumor collection, tumor differentiation, smoking and alcohol history, initial treatment modalities, postoperative radiotherapy and/or chemotherapy, and follow-up data including recurrence and survival. Time points were calculated from the start date of treatment, and recurrence (a new tumor event) and survival (time to death from SNSCC or other causes or the last follow-up) were also assessed.

### Bulk RNA sequencing analysis

RNA was extracted from tissue specimens in the accordance with the RNeasy® Mini kit (Qiagen, Cat# 74104) following the manufacturer’s instructions. RNA concentration was quantified by measuring UV absorbance at 260 nm with spectrophotometer and RNA integrity was checked using an Agilent Technologies 2100 Bioanalyzer with an RNA Integrity Number (RIN) value greater than or equal to 7.0. Following RNA extraction, sequencing libraries were constructed using the Illumina TruSeq RNA Library Prep Kit (Illumina, Cat#20020189) following the manufacturer’s guidelines. The raw data were processed by trimming with Trim Galore, and the reads were subsequently aligned to the human reference genome hg38 using the STAR aligner^71^ with default parameters. Gene expression levels in each sample were normalized using fragments per kilobase of transcript per million mapped reads (FPKM). Differentially expression analysis was performed with DESeq2 v1.34.0^72^, identifying significantly upregulated or downregulated DEGs in SNSCC tumors compared with normal tissues based on FPKM > 2, log2 fold-change > 1, and adjusted *p*-value < 0.05. Functional enrichment of DEGs was performed using Metascape^73^ (https://metascape.org), and GSEA was conducted using the hallmark gene set from the Molecular Signatures Database (MsigDB)^74, 75^ (https://www.gsea-msigdb.org/gsea/index.jsp). The PROGENy package v1.16.0^76^ was employed to evaluate the activity of 14 key signaling pathways in each sample.

### Deconvolution of bulk RNA-seq data

The CIBERSORTx^77^ (https://cibersortx.stanford.edu) deconvolution algorithm was utilized to estimate the relative abundances of 22 immune cell types (LM22) using gene expression levels from bulk RNA-seq performed on tumor and normal tissues of patients with SNSCC.

### DNA methylation EPIC array analysis

DNA was extracted from tissue specimens in the accordance with the QIAamp DNA Mini (Qiagen, Cat# 51304) following the manufacturer’s instructions. DNA methylation profiles were generated using the Illumina HumanMethylation EPIC array platform. These profiles were subsequently integrated with publicly accessible DNA methylation EPIC array data of SNSCC tumor and normal samples from the GSE196228 dataset^22^. All DNA methylation data were processed and analyzed using the ChAMP package v2.24.0^78^ in the R environment. Following a pipeline that included filtering of unreliable detections and probes, quality control, normalization, and batch effect removal via the champ.runCombat function, we proceeded to identify DMPs. The comparison between SNSCC tumor and normal samples was conducted using the champ.DMP function. Significant DMPs were defined by stringent criteria, which included an adjusted *p*-value < 0.05 and a beta-value difference exceeding 20% (β-value > 0.2). Motif enrichment analysis of differentially methylated CpG sites was conducted with the HOMER software v4.11.1^79^ utilizing the findMotifsGenome.pl function, with the parameters set to “size 200 -mask”.

### Nuclei isolation from human tissues and snRNA/ATAC-seq library preparation

Nuclei were isolated from solid tissues using a Singulator 100 system (S2 Genomics) following the manufacturer’s protocol. Debris was removed using a Percoll gradient. Nuclei concentration was determined using a LUNA-FL™ Automated Fluorescence Cell Counter (Logos Biosystems), and morphology was examined by microscopy. The isolated nuclei were then incubated in a Transposition Mix containing Transposase.

Libraries were prepared using the Chromium controller (10x Genomics) according to the Chromium Single Cell Multiome ATAC + Gene Expression protocol (CG000338). Briefly, transposed nuclei were mixed with master mix and loaded with Single Cell Multiome gel beads and partitioning oil into a Chromium Chip J. The gel beads contained poly (dT) sequences for gene expression (GEX) library preparation and spacer sequences for ATAC library preparation.

Gel bead-in-emulsion (GEM) incubation resulted in 10x barcoded DNA from transposed DNA (for ATAC) and 10x barcoded cDNA from polyadenylated mRNA (for GEX). After pre-amplification, the PCR product was divided for separate ATAC and GEX library construction steps. For the ATAC library, P7 adapters and sample indices were added to pre-amplified transposed DNA during library construction via PCR. For the GEX library, pre-amplified cDNA underwent PCR amplification, followed by fragmentation, end repair, A-tailing, adapter ligation, and index PCR.

### Library quantification and snRNA/ATAC-seq

The purified libraries were quantified using qPCR according to the qPCR Quantification Protocol Guide (KAPA Biosystems). Library quality was assessed using an Agilent Technologies 4200 TapeStation (Agilent technologies). Sequencing was performed on a HiSeq platform (Illumina) following the read length in the user guide.

### snRNA/ATAC-seq data pre-processing and quality control

Raw sequencing data (FASTQ files) were aligned to the hg38 reference genome and quantified using Cell Ranger ARC v2.0.2. cellranger-arc count pipeline produced barcoded count matrices for gene expression and ATAC data. These matrices and fragment files were then processed using Seurat v4.3.0^80^ and Signac v1.9.0^81^ software packages.

SoupX v1.6.2^82^ was employed to assess and mitigate background RNA contamination. The autoEstCont function was used to estimate contamination fractions, which were manually adjusted to 0.2 for tumor samples using setContaminationFraction. Background noise was subsequently eliminated using the adjustCounts function with default parameters. To generate a unified peak set from the snATAC-seq data of all samples, we used the reduce function from GenomicRanges v1.46.1^83^. These combined peaks were then quantified across all datasets. The scDblFinder v1.8.0^84^ was employed to identify and remove potential doublets. We implemented stringent quality control criteria, retaining cells with nCount_RNA > 1000, nCount_ATAC > 1000, percent.mt < 3, and TSS.enrichment > 1.5. Sample-specific thresholds were also implemented for each sample as follows: HC1: nCount_ATAC < 20000 & nCount_RNA < 20000 & nucleosome_signal < 0.7 HC2: nCount_ATAC < 25000 & nCount_RNA < 20000 & nucleosome_signal < 0.7 CA5: nCount_ATAC < 20000 & nCount_RNA < 20000 & nucleosome_signal < 1.5 CA8: nCount_ATAC < 5000 & nCount_RNA < 15000 & nucleosome_signal < 1

### Joint snRNA/ATAC-seq data workflow

Post-quality control, snRNA-seq data were processed using Seurat package. Normalization, variable feature identification (top 3000 genes), scaling, and principal component (PC) analysis were performed. Batch effects were removed using Harmony v0.1.1^85^. RunUMAP function was applied to the first 30 Harmony-corrected PC dimensions for RNA data visualization.

snATAC-seq peak calling was conducted using MACS2 v2.2.6^86^ via the Signac package. Peaks overlapping with hg38 blacklist regions were excluded. The resulting peak set was used to create a new chromatin assay within the Seurat object. Top features were identified (min.cutoff = 5), followed by TF-IDF normalization and partial singular value decomposition. Batch effects of ATAC datasets were addressed using the RunHarmony function in Harmony package on latent semantic indexing (LSI) components. Uniform manifold approximation and projection (UMAP) visualization was employed the RunUMAP function based on the second through 30th Harmony-corrected LSI components.

Weighted nearest neighbor (WNN) analysis was used to integrate gene expression and chromatin accessibility data. The FindMultiModalNeighbors function combined gene expression (PCA) and ATAC (LSI) data. WNN-derived clusters were visualized using UMAP projection with RunUMAP function, setting nn.name to weighted.nn. Clustering was performed using the FindClusters function with a resolution of 0.2 and algorithm 3, resulting in 28 clusters. Low-quality cells exhibiting low gene detection with high mitochondrial reads and potential doublets showing double-positive cell type marker gene expression, were excluded from downstream analysis.

### Differential expression analysis in snRNA/ATAC-seq data

Differential expression analysis was performed using Seurat v4.3.0. DEGs in major cell types were identified using the FindAllMarkers function (parameters: only.pos = TRUE, min.pct = 0.1, logfc.threshod =1). Across malignant cell subpopulations, the logfc.threshold was adjusted to 0.58. DEGs between endothelial cell subclusters were determined using the FindMarkers function (min.pct = 0.1, logfcthreshold = 1). DARs between each cell type within tumor and HC tissues were identified using the FindMarkers function with min.pct = 0.02, test.use = LR, latent.vars = nCount_peaks, and logfc.threshold = 1. Genomic distribution of DARs was analyzed using HOMER v4.11.1 annotatePeaks.pl script with human genome hg38 as reference. All significant DEGs and DARs were filtered with adjusted *p*-value < 0.05.

### Functional analysis in snRNA/ATAC-seq data

GO analysis of DEGs was performed using Metascape. Functional annotation of DARs was conducted using annotatePeaks.pl. with hg38 and -go options in Homer software. The PROGENy package v1.16.0^76^ was employed to evaluate the activity of 14 key signaling pathways in snRNA/ATAC-seq datasets.

### Gene-peak linkage analysis

Gene-peak linkages were evaluated using the LinkPeaks function in Signac. Significant linked peaks (*p*-value < 0.05 and linkage score > 0) were integrated with DARs in epithelial cells, fibroblasts, and endothelial cells for finding linked DARs with genes. Further overlapping with DEGs in each tumor-enriched cell type compared with their normal compartments using FindMarkers function with min.pct = 0.1, logfc.threshold = 1, and adjusted *p*-value below 0.05, were found DEGs-linked DARs in each cell type.

### Transcription factor analysis using chromvar

TF binding motifs were annotated using the JASPAR 2020 database^87^ and analyzed with chromVAR v1.16.0^88^ to estimate TF activity. These values were used to generate the motif heatmaps and feature plots. Differential motif activity was accessed using FindAllMarkers function with only.pos = TRUE, min.fxn = rowMeans, fc.name = avg_diff, logfc.threshold = 0.58, and adjusted *p*-value < 0.05.

### InferCNV

CNV within epithelial cell subpopulations were defined using the InferCNV algorithm v1.10.1 (https://github.com/broadinstitute/infercnv). Raw count data, extracted from RNA assay in the seurat object of epithelial cell clusters, were utilized to generate InferCNV object via CreateInfercnvObject function. Ciliated, mucous, and serous cells served as the reference group for this analysis. To detect CNV signals, we employed the infercnv::run function, setting the cutoff for the minimum average read count per gene to 0.1, as recommended for 10x Genomics data. Subsequently, the derived CNV data were integrated into the seurat object using the infercnv::add_to_seurat function to facilitate visualization and downstream analyses.

### Module score

Module score for specific gene sets were calculated using the AddModuleScore function in Seurat. Signature gene lists were derived from the top 100 upregulated and downregulated DEGs from bulk RNA-seq analysis, the HALLMARK gene sets for hypoxia and glycolysis from the MSigDB, and angiogenesis signature genes from previous studies^89, 90^.

### Gene set variation analysis

Gene set variation analysis (GSVA) v1.42.0^91^ was used to generate signature scores for each TC subset (TC1-TC5) in HNSC patients from TCGA data. These scores were used for overall survival analysis of TC subpopulations.

### Cell-cell integration analysis

Intercellular signaling networks were analyzed using CellChat v1.6.1^92^. We selected the secreted signaling pathways from the CellChat human database. The CellChatDB database was supplemented with ADM-RAMP2 and ADM-RAMP3 ligand-receptor pairs to dissect adrenomedullin signaling pathways. CellChat objects were created using the createCellChat function. Ligand-receptor interactions between cell populations were determined using the CellChat pipeline with default parameters. For comparative analysis, independent CellChat objects were generated for SNSCC and HC tissues, then integrated using the mergeCellChat function. Differential interaction analysis was performed to identify significant changes in signaling pathways and ligand-receptor pairs between SNSCC and HC tissues. Communication networks were visualized using stacked bar, bubble, and circle plots.

### TCGA-HNSC analysis

Gene expression matrix and clinical information, including stage and survival rate of HNSC patients, were obtained from the UCSC Xena web portal (https://xenabrowser.net/datapages/). We generated correlation analysis and overall survival curves, excluding patients with missing stage or survival period information. ADM gene expression and signature scores of each TC subpopulation were used to divide patients into two groups at the 50th percentile. *P* values were calculated using log-rank test.

### Statistical analysis

Statistical analyses were conducted using R software (versions 4.1.0 and 4.1.1) and GraphPad Prism 9. Results are presented as mean ± standard error of the mean (SEM) unless otherwise specified. Statistical significance was defined as *p*-value < 0.05, as determined by two-tailed unpaired t-tests or two-way ANOVA where appropriate. In box plots, center lines show medians, box limits indicate 25th and 75th percentiles, and whiskers extend to minimum and maximum values or 10th and 90th percentiles, as specified in figure legends. Correlations between variables were assessed using Pearson’s correlation coefficient. Statistical significance of correlations was evaluated using two-tailed tests, with *p*-value < 0.05 considered significant. Survival analyses were performed using the Kaplan-Meier method. Differences in survival curves were assessed using the log-rank test. Patients were stratified into high and low groups based on the median expression level of genes of interest or signature scores.

## Data Availability

The datasets generated in this study are available in the GEO repository under SuperSeries accession GSE278149. This SuperSeries is composed of the following SubSeries: GSE278145 (bulk RNA-seq), GSE278138 (DNA methylation EPIC array), and GSE277797 (snRNA/ATAC-seq). The public dataset analyzed in this study is available under the GEO accession GSE196228^22^. The HNSC dataset from TCGA was obtained from the UCSC Xena web portal (https://xenabrowser.net/datapages/).

## Code Availability

This study did not generate new code or unique materials. All analyses were performed using existing software as detailed in the Methods. For additional information required to replicate the analyses in this manuscript, please contact the corresponding authors.

## Acknowledgements

This study was supported by a grant (2023IL0014 and 2023IP0057) from the Asan Institute for Life Sciences, Asan Medical Center, Seoul, Korea and the National Research Foundation (NRF), Republic of Korea, grant (RS-2024-00340411 and NRF-2022R1A2C1011187), which is funded by the Ministry of Science and ICT (MSIT)

## Author contributions

C.Y. conceptualized, designed, and performed most of the experiments and conducted bioinformatic analyses. Jaewoo P., J.Y.J., M.S.Y., and Y.S.C. contributed to the experiments. Keunsoo K. and Jihwan P. contributed to bioinformatic analyses and provided expertise. Kyuho K. and J.H.K. conceptualized and supervised the study and edited the manuscript. All authors reviewed and provided input on the manuscript.

## Competing interests

The authors declare no competing interests.

## Additional information

**Supplementary information:** Figure S1 - S7.

## Notes

### Competing Interest Statement

The authors have declared no competing interest.

