## Supplementary figures for "Multi-omic profiling identifies malignant subpopulations and a hypoxia-driven angiogenic axis as therapeutic vulnerabilities in sinonasal squamous cell carcinoma"

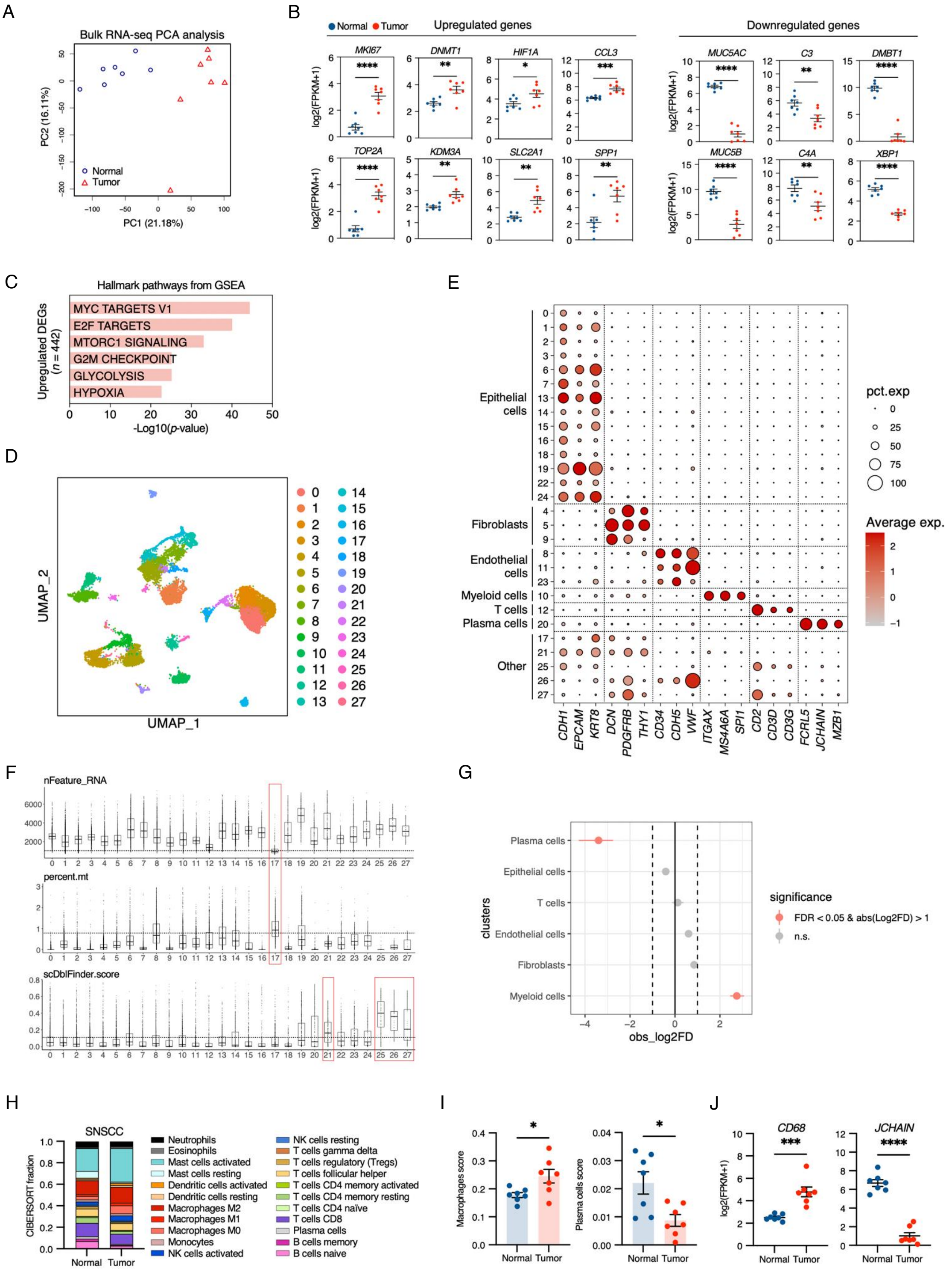

**Supplementary Figure S1. Transcriptomic and cellular landscape of SNSCC.**

(**A**) PCA plot of bulk RNA-seq data of tumor and normal samples of SNSCC patients.

(**B**) Relative expression of select genes associated with representative GO terms of

SNSCC up- or downregulated gene sets from bulk RNA-seq data identified in Fig. 1C. Each dot represents one donor. Data are shown as mean ± SEM. *P* values were determined by two-tailed unpaired t-test. **p* < 0.05, ***p* < 0.01, ****p* < 0.001, *****p* < 0.0001. (**C**) The top six GSEA hallmark pathways enriched in SNSCC tumor compared to normal samples. (**D**) UMAP visualization of the 18,319 cells computed using WNN profiles, colored by 28 cell clusters. (**E**) Dot plot showing gene expression of known markers in 28 cell clusters. Annotated cell types with their corresponding cell clusters are listed on the left side. (**F**) Violin plots displaying the number of genes (nFeature_RNA), percentage of mitochondrial genes (percent.mt), and doublet score calculated using scDblFinder (scDblFinder.score) in each cell cluster. (**G**) Differences in single-cell proportions of six major cell types between SNSCC tumor and HC tissues. Dashed lines indicate the log2 fold change of 1. A significant difference in cell proportion is determined by false discovery rate (FDR) less than 0.05. (**H**) CIBERSORT analysis of 22 infiltrating immune cell types in SNSCC tumor and normal tissues summarized from calculated mean values for each sample (100 permutations). (**I**) Differences in macrophage (M0 + M1 + M2) and plasma cell scores between SNSCC tumor and normal tissues. Each dot represents one donor. Data are shown as mean ± SEM. *P* values were determined by two-tailed unpaired t-test. **p* < 0.05, ***p* < 0.01, ****p* < 0.001, *****p* < 0.0001. (**J**) Gene expression levels of *CD68* and *JCHAIN* in SNSCC tumor versus normal samples from bulk RNA-seq. Each dot represents one donor. Data are shown as mean ± SEM. *P* values were determined by two-tailed unpaired t-test. **p* < 0.05, ***p* < 0.01, ****p* < 0.001, *****p* < 0.0001.

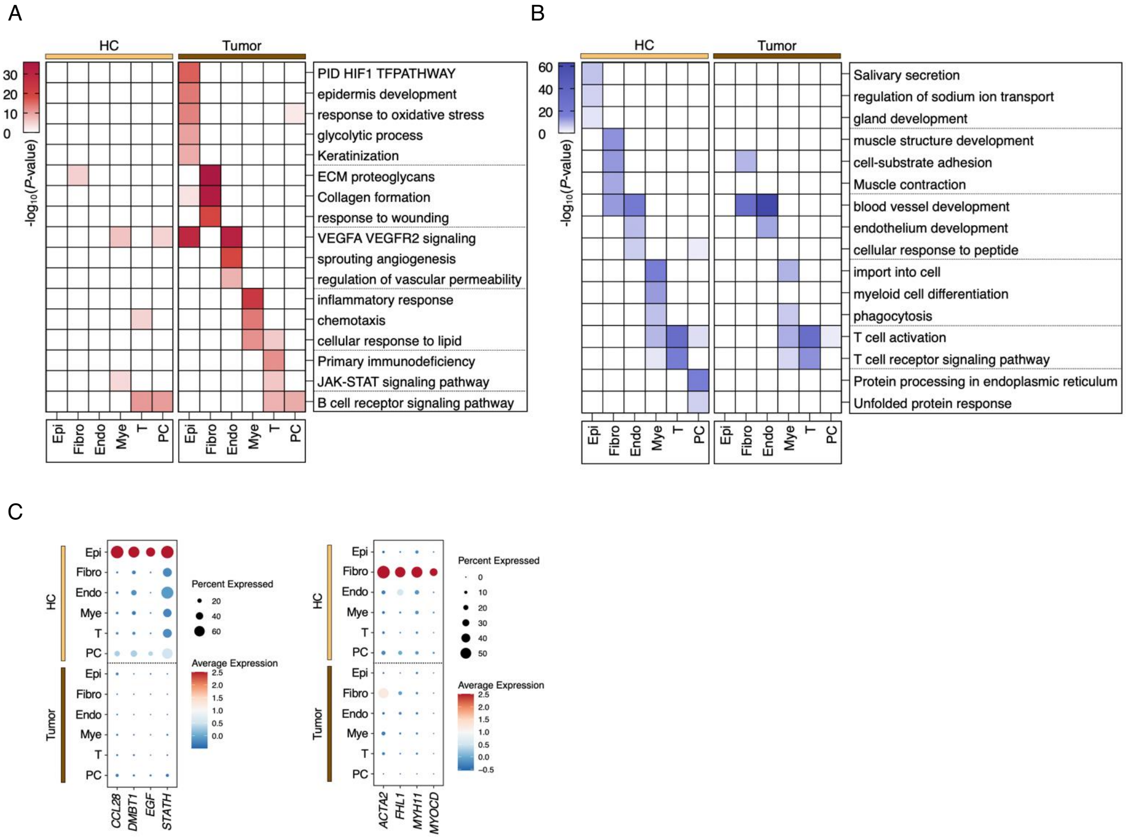


**Supplementary Figure S2. GO analysis of cell type-specific markers in SNSCC and HC tissues.**

(**A**) Heatmap representing the *p*-value significance of enriched GO terms for marker genes in each cell type within SNSCC tumor tissues. (**B**) Heatmap representing the *p*-value significance of enriched GO terms for marker genes in each cell type within HC tissues. (**C**) Dot plots showing expression of select marker genes associated with representative GO terms in epithelial cells (left) and fibroblasts (right) within HC tissues.

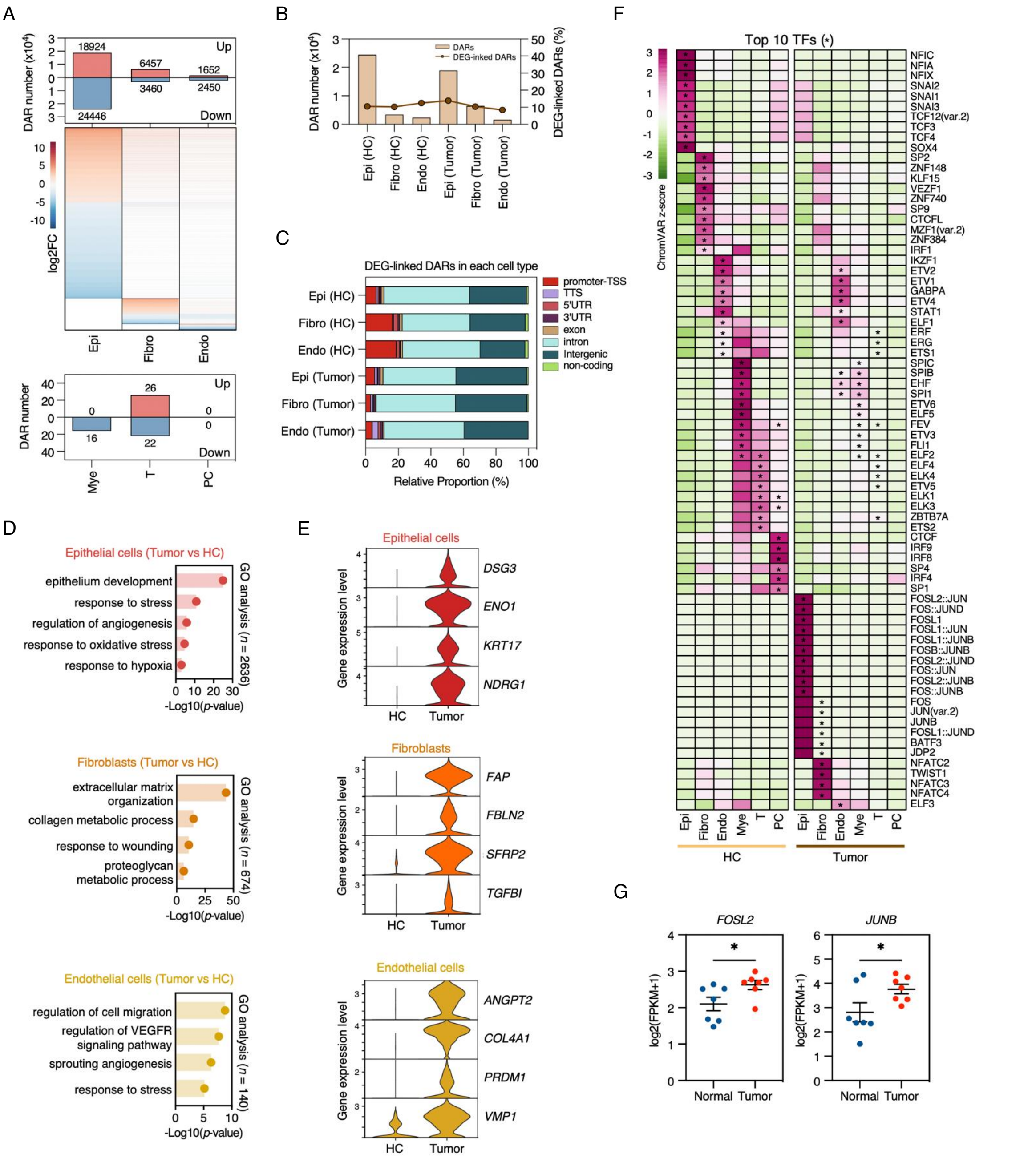

**Supplementary Figure S3. Differential chromatin accessibility and TF activity in SNSCC.**

(**A**) The number (top) and log2 fold change (middle) of up- and downregulated DARs in epithelial cells, fibroblasts, and endothelial cells within SNSCC tumors compared with HC samples. The number of up- and downregulated DARs in myeloid, t, and plasma cells within SNSCC tumors compared to HC samples (bottom). (**B**) Bar plot displaying DAR numbers with proportion of DEG-linked DARs in each epithelial cells, fibroblasts, and endothelial cells within SNSCC tumor and HC samples. (**C**) Genomic distribution of annotated DEG-linked DARs in epithelial cells, fibroblasts, and endothelial cells within SNSCC tumors and HC samples, calculated by HOMER. (**D**) GO analysis of upregulated DEG-linked opened DARs in epithelial cells (top), fibroblasts (middle), and endothelial cells (bottom) within SNSCC tumors identified in Fig. 3A, performed by HOMER. (**E**) Violin plots showing gene expression levels of selected opened DAR-linked upregulated DEGs in epithelial cells (top), fibroblasts (middle), and endothelial cells (bottom) within SNSCC tumors identified in Fig. 3A. (**F**) Heatmap showing chromVAR motif activity of top 10 enriched TF binding motifs in each cell type within SNSCC tumor and normal samples. (**G**) Expression levels of *FOSL2* and *JUNB* in SNSCC tumor versus normal samples from bulk RNA-seq. Each dot represents one donor. Data are shown as mean ± SEM. *P* values were determined by two-tailed unpaired t-test. **p* < 0.05, ***p* < 0.01, ****p* < 0.001, *****p* < 0.0001.

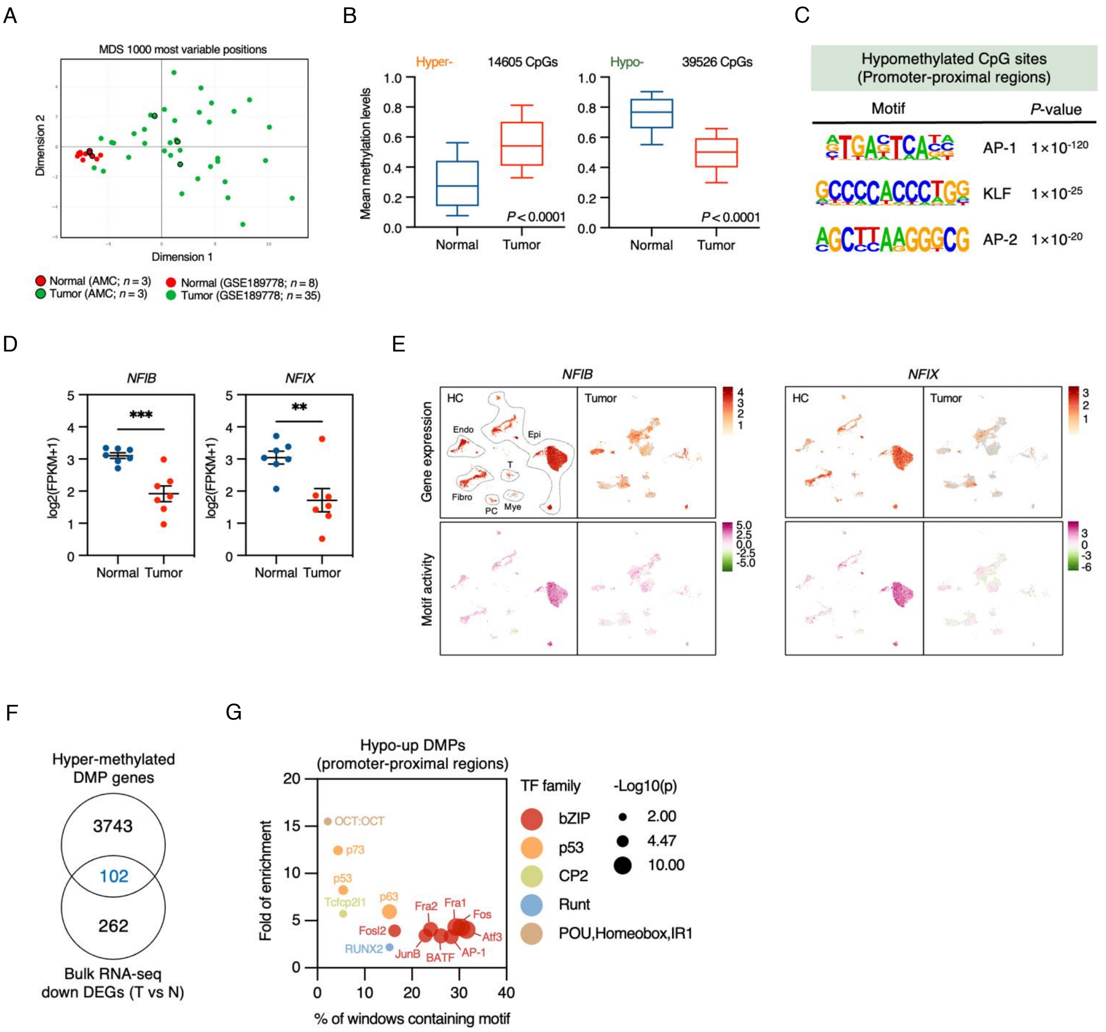


**Supplementary Figure S4. DNA methylation patterns and TF activity in SNSCC.**

(**A**) MDS plot of EPIC array data for SNSCC tumor and normal tissues from our data and GSE189778. (**B**) Mean methylation levels of hypo- and hypermethylated DMPs in SNSCC tumor and normal samples. Boxes encompass the 25th to 75th percentile, Whiskers extend to the 10th and 90th percentiles, and center line indicates the median. *P* values were determined by two-tailed unpaired t-test. (**C**) Significantly enriched binding motifs in hypomethylated DMPs in promoter-proximal regions revealed by de novo motif analysis using HOMER software. (**D**) Expression levels of *NFIB* and *NFIX* in SNSCC tumor versus normal samples from bulk RNA-seq. Each dot represents one donor. Data are shown as mean ± SEM. *P* values were determined by two-tailed unpaired t-test. *$p < 0.05$, **$p < 0.01$, ***$p < 0.001$, ****$p < 0.0001$. (**E**) UMAP plots for gene expression (upper) and chromVAR motif activity (lower) of NFIB and NFIX split by tissue type. (**F**) Venn diagram showing the number of hypermethylated downregulated (hyper-down) genes in SNSCC tumor compared with normal samples. (**G**) Bubble plot showing enriched TF motifs in hypo-up DMPs within promoter-proximal regions of SNSCC tumors compared to normal samples, analyzed using HOMER software. Color range represents different TF families and circle size corresponds to *p*-value.

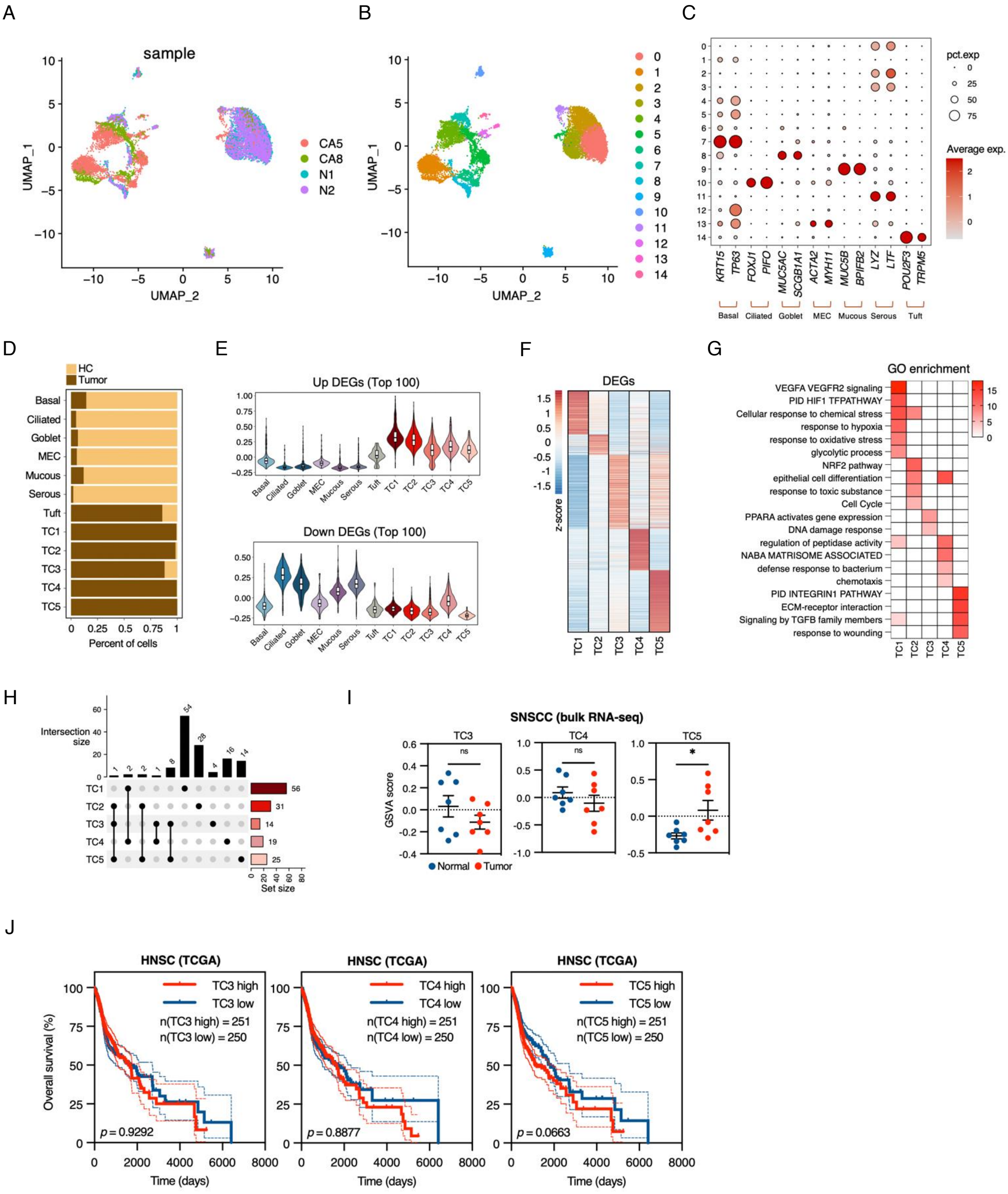


**Supplementary Figure S5. Characterization of epithelial cell populations in SNSCC.**

(**A**) UMAP plot of epithelial compartments in SNSCC tumor and HC samples computed by WNN, colored by sample. (**B**) UMAP plot of epithelial compartments in SNSCC tumor and HC samples computed by WNN, colored by 15 cell clusters. (**C**) Dot plot showing gene expression of cell type marker genes in 15 cell clusters. (**D**) Cell proportions of SNSCC tumor and HC tissues in each epithelial population. (**E**) Violin plots representing signature scores of top 100 up- (upper) and downregulated (lower) DEGs from bulk RNA-seq data in all epithelial cell populations. (**F**) Heatmap of DEGs across malignant cell subpopulations. (**G**) Heatmap representing the *p*-value significance of enriched GO terms for DEGs across malignant cell types. (**H**) Upset plot showing the number of integrated genes of DEGs across malignant cell types and upregulated DEGs in SNSCC tumors compared to normal samples from bulk RNA-seq data. (**I**) Signature scores of TC3, TC4, and TC5 in SNSCC tumor versus normal samples from bulk RNA-seq. Each dot represents one donor. Data are shown as mean ± SEM. *P* values were determined by two-tailed unpaired t-test. $^{*}p < 0.05$, $^{**}p < 0.01$, $^{***}p < 0.001$, $^{****}p < 0.0001$, ns: not significant. (**J**) Kaplan-Meier curves of overall survival according to high and low groups in HNSC patients from TCGA dataset, stratified by TC3 (left), TC4 (middle), and TC5 (right) signature scores (50th percentile). *P* values were determined by log-rank test.

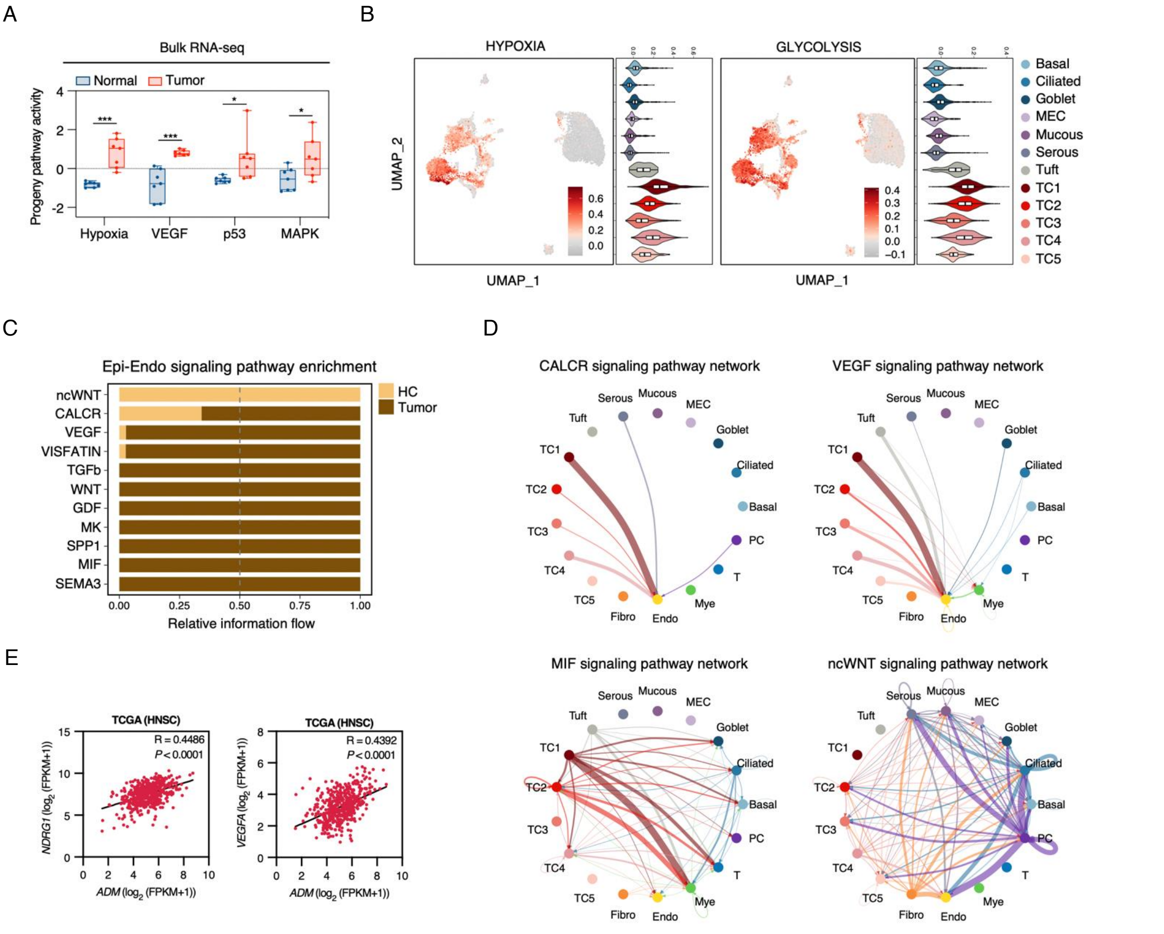


**Supplementary Figure S6. Signaling pathway analysis and cell-cell communication in SNSCC.**

(**A**) Box plots showing the activity of signaling pathways that significantly upregulated in SNSCC tumors compared to normal tissues, determined using PROGENy. (**B**) UMAP and violin plots of hypoxia (left) and glycolysis (right) signature scores in epithelial cells. (**C**) Stacked bar plot displaying enriched signaling pathways from epithelial cells to endothelial cells between SNSCC tumor and HC samples. (**D**) Circos plots of CALCR, MIF, VEGF, and ncWNT signaling pathways between all epithelial and endothelial states. The thickness of edge represents the communication probability. (**E**) Pearson correlations between the gene expression levels of *ADM* and

*NDRG1* (left) or *ADM* and *VEGFA* (right) from TCGA-HNSC dataset.

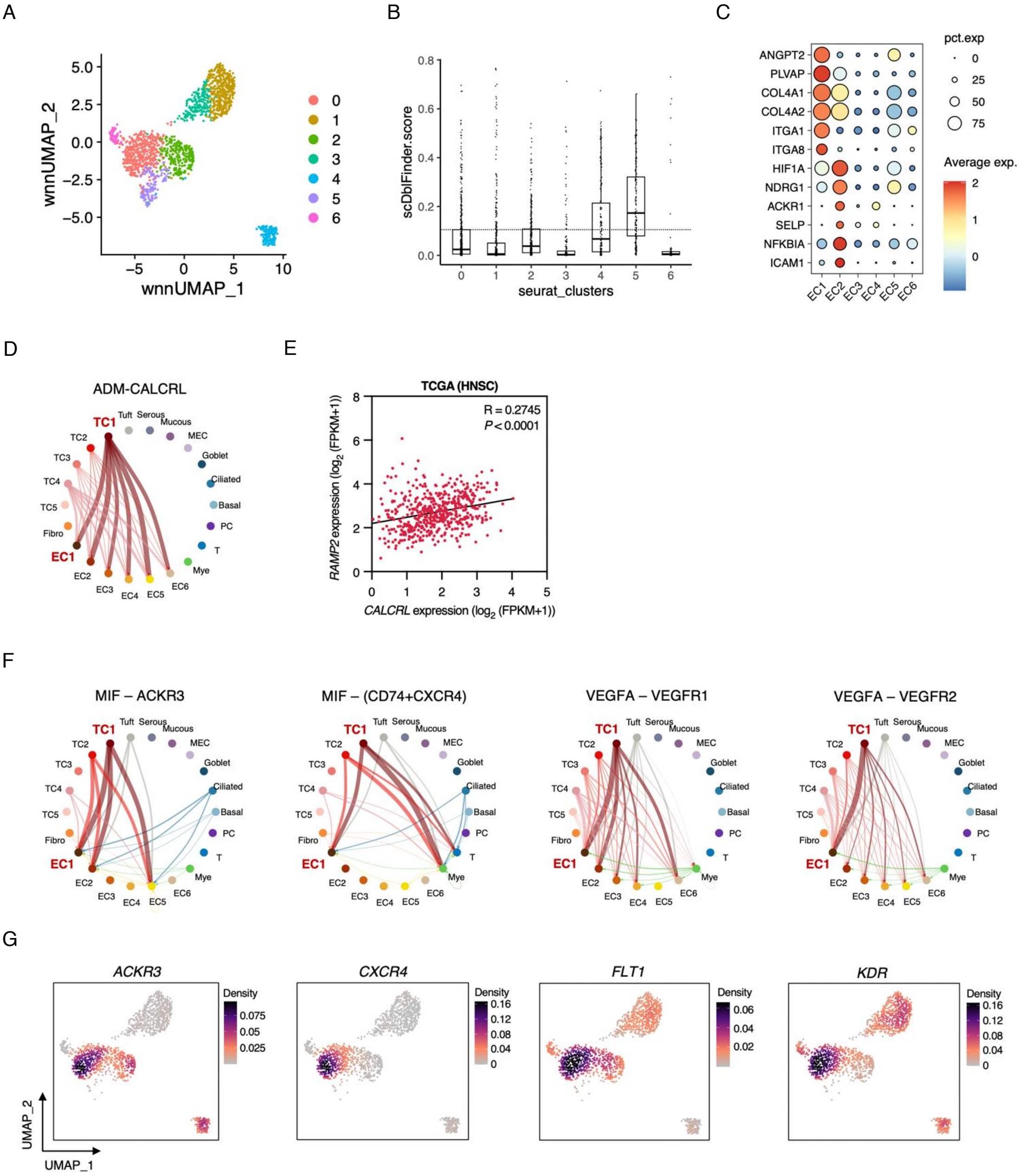


**Supplementary Figure S7. Analysis of endothelial cell populations and their interactions in SNSCC.**

(**A**) UMAP visualization of endothelial compartments in SNSCC tumor and HC samples by WNN analysis, colored by 7 cell clusters. (**B**) Box plots displaying doublet scores calculated using scDblFinder (scDblFinder.score) in each endothelial cell subcluster. (**C**) Dot plot of expression levels of select upregulated DEGs in EC1 and

EC2 compared to other endothelial cell subclusters identified in Fig. 7D. (**D**) Circos plot of inferred ADM-CALCRL interactions between all epithelial and endothelial states. The thickness of edge represents the communication probability. (**E**) Pearson correlations between the gene expression levels of *CALCRL* and *RAMP2* from TCGA-HNSC dataset. (**F**) Circos plots of inferred MIF-ACKR3, MIF-(CD74+CXCR4), VEGFA-VEGFR1, and VEGFA-VEGFR2 interactions between all cell types with epithelial and endothelial subpopulations. The thickness of edge represents the communication probability. (**G**) Density plot of *ACKR3*, *CXCR4*, *FLT1*, and *KDR* expression in endothelial compartments.
